# Dual RNA-seq provides insight into the biology of the neglected intracellular human pathogen *Orientia tsutsugamushi*

**DOI:** 10.1101/743641

**Authors:** Bozena Mika-Gospodorz, Suparat Giengkam, Alexander J. Westermann, Jantana Wongsantichon, Willow Kion-Crosby, Suthida Chuenklin, Loo Chien Wang, Piyanate Sunyakumthorn, Radoslaw M. Sobota, Selvakumar Subbian, Jörg Vogel, Lars Barquist, Jeanne Salje

**Affiliations:** Helmholtz Institute for RNA-based Infection Research (HIRI), Helmholtz Centre for Infection Research (HZI), Würzburg, Germany; Mahidol-Oxford Tropical Medicine Research Unit, Faculty of Tropical Medicine, Mahidol University, Bangkok, Thailand; Institute for Molecular Infection Biology (IMIB), University of Würzburg, Würzburg, Germany; Rutgers, the State Univeristy of New Jersey, New Jersey, USA; Functional Proteomics Laboratory, Institute of Molecular and Cell Biology, Agency for Science, Technology and Research (A*STAR), Singapore; Armed Forces Research Institute of Medical Sciences, Bangkok, Thailand; Faculty of Medicine, University of Würzburg, Würzburg, Germany; Public Health Research Institute, Rutgers University, New Jersey, USA; Centre for Tropical Medicine and Global Health, Nuffield Department of Medicine, University of Oxford, Oxford, United Kingdom; SingMass - National Mass Spectrometry Laboratory, Institute of Molecular and Cell Biology, Agency for Science, Technology and Research (A*STAR), Singapore

**Keywords:** neglected and emerging pathogens, intracellular bacteria, dual RNA-seq, transcriptomics, host-pathogen cell biology, bacterial virulence, antisense transcription

## Abstract

Emerging and neglected diseases pose challenges as their biology is frequently poorly understood, and genetic tools often do not exist to manipulate the responsible pathogen. Organism agnostic sequencing technologies offer a promising approach to understand the molecular processes underlying these diseases. Here we apply dual RNA-seq to *Orientia tsutsugamushi* (Ot), an obligate intracellular bacterium and the causative agent of the vector-borne human disease scrub typhus. Half the Ot genome is composed of repetitive DNA, and there is minimal collinearity in gene order between strains. Integrating RNA-seq, comparative genomics, proteomics, and machine learning, we investigated the transcriptional architecture of Ot, including operon structure and non-coding RNAs, and found evidence for wide-spread post-transcriptional antisense regulation. We compared the host response to two clinical isolates and identified distinct immune response networks that are up-regulated in response to each strain, leading to predictions of relative virulence which were confirmed in a mouse infection model. Thus, dual RNA-seq can provide insight into the biology and host-pathogen interactions of a poorly characterized and genetically intractable organism such as Ot.

## Introduction

Improved surveillance and diagnostics have led to the recognition of previously neglected bacteria as serious pathogens, whilst human population growth, globalization and increased travel have contributed to the emergence of new pathogens and changing patterns of infectious disease. The biology of neglected and emerging pathogens is often poorly understood, but is essential to developing therapeutic and preventative strategies. Obligate intracellular pathogens present additional challenges, as many cause diseases that are difficult to diagnose and are difficult to manipulate experimentally.

Obligate intracellular bacteria include the Rickettsiales, an order which includes the arthropod and nematode symbiont *Wolbachia* as well as a number of human and veterinary pathogens. *Orientia tsutsugamushi* (Ot, Class Alphaproteobacteria, Order Rickettsiales, Family Rickettsiaceae) causes the mite-borne human disease scrub typhus, a leading cause of severe febrile illness in the Asia Pacific region^1^, home to roughly two thirds of the world’s population. Locally acquired cases in the Middle East and Latin America suggest that this disease may be more widespread than previously appreciated^2, 3^. Under-recognition and under-reporting are a major problem in scrub typhus because unambiguous diagnosis is difficult, and awareness is low amongst many clinicians. Symptoms are non-specific and include headache, fever, rash, and lymphadenopathy beginning 7-14 days after inoculation via a feeding larval stage mite. If untreated, this can progress to cause multiple organ failure and death. In the mite vector Ot colonizes the ovaries and salivary glands. During acute infection of its mammalian host, the bacteria infect endothelial cells, dendritic cells and monocytes/macrophages at the mite bite site^4^, and then disseminate via blood and lymphatic vessels to multiple organs including lung, liver, kidney, spleen and brain^5^.

Ot strains are highly variable in terms of antigenicity and virulence. Hundreds of strains have been described based on differences in the sequence of the surface protein TSA56^9–11^. These strains are classified into 7 geographically diverse serotype groups, dominated by the Karp, Kato and Gilliam groups^12^. As with many other pathogenic bacteria, whole genome sequencing has revealed that serotype-based groupings do not necessarily reflect phylogenetic relationships^8^. Different strains of Ot exhibit different levels of virulence^19–21^, dependent on both bacterial and host genotype. For example, strain Karp (serotype group Karp) causes lethal infection in BALBc and C3H/He mice at low doses, strain Gilliam (serotype group Gilliam) causes lethal infection in C3H/He but not BALBc mice at similar doses, whilst strain TA716 (serotype group TA716) does not cause lethal infection in either mouse model at similar doses^20, 22^. The underlying causes of this variation in infection outcomes remain obscure.

Dual RNA-seq quantifies RNA transcripts of intracellular pathogens and host cells in a single experiment^24, 25^, and can provide insight into both the host and pathogen response to infection. For example, dual RNA-seq has been used to study obligate intracellular *Chlamydia trachomatis*^26^ revealing the rewiring of *Chlamydia* metabolism during the onset of an infection of human epithelial cells, together with the corresponding host responses. Here we apply dual RNA-seq to deepen our understanding of the RNA biology of Ot and its consequences for virulence. We survey the transcriptome of Ot strain Karp, identifying non-coding RNAs and transcribed operons in a genome broken by frequent recombination and transposition of the rickettsial <u>a</u>mplified <u>g</u>enetic <u>e</u>lement (RAGE) integrative and conjugative element (ICE)^6, 7^. Integrating proteomic measurements, we further provide evidence that RAGE genes are regulated through prevalent antisense transcription. Finally, we compare infection between strain Karp and strain UT176 identifying a core host response to Ot dominated by type-I interferon signaling, as well as distinct immune responses to each strain. We show that this in turn leads to different outcomes in a mouse model of scrub typhus.

## Results

### Dual RNA-seq of *Orientia tsutsugamushi* infecting endothelial cells

We focused on two Ot clinical isolates: Karp, taken from a patient in New Guinea in 1943^27^, and UT176, a separate strain in the same serotype group as Karp, taken from a patient in northern Thailand in 2004^28^. These strains are closely related, with a sequence identity of 95% in their TSA56 gene (used to classify strains)^8^. Consistent with a closed pan-genome for Ot, the gene content of Karp and UT176 are similar, with differences primarily in gene copy number, pseudogenes, and gene order along the genome. Human vein endothelial cells (HUVEC) were selected as host cells due to their similarity to cell types involved in both early and advanced infection. HUVEC cells were infected with bacteria at an MOI of 100:1 and grown for 5 days (Fig. 1A), by which point host cells were heavily loaded with bacteria (Fig. 1C, Supp. Figs 1, 2). Uninfected HUVEC cells were grown in parallel. After 5 days total RNA was isolated, depleted for rRNA, converted to cDNA and sequenced to ∼35 million reads per library using Illumina technology. Reads were mapped to the completed genomes of Karp, UT176^8^ and, in parallel, the human genome. As the *Orientia* genome is repeat-rich, we additionally applied model-based quantification with Salmon^29^ which uses uniquely mapping reads to assign multi-mapping reads to these transcriptomes to improve our estimates of transcript abundance (see Methods).

**Figure 1.**
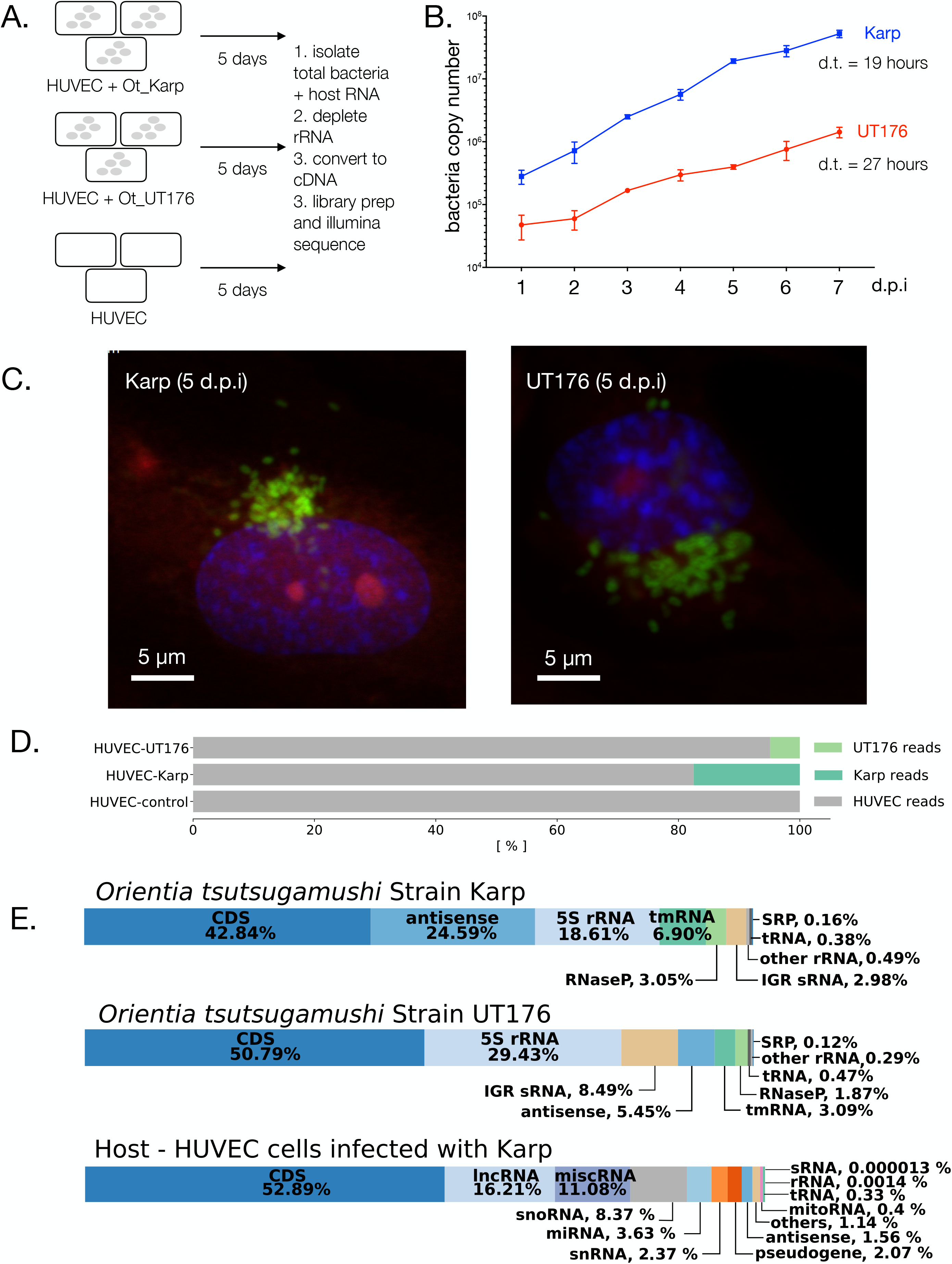
Overview. **A.** Schematic experimental overview. HUVEC = human umbilical vein endothelial cell. **B.** Growth curve showing replication of Ot in cultured HUVEC cells. Bacteria were grown in 24 well plates and the total bacteria per well is shown. Mean and SD from three independent replicates are shown. **C.** Confocal microscopy images of Ot in HUVEC cells 5 days post infection. Additional images and time points are shown in Supp. Fig. 1 and 2. Blue = DAPI (DNA), Red = Evans blue (host cells), green = Ot labelled with Alexa488-click-methionine. **D.** RNA mapping statistics showing the fraction of host and Ot RNA for each condition. The first replicate of the experiment is shown. Individual results for each replicate are shown in Supp. Fig. 3. **E.** Percentage of RNA-seq reads assigned to different classes of RNA in Karp, UT176 and HUVEC.

We observed 17.1-17.5% bacterial reads in HUVECs infected with Karp and 2.8-4.9% bacterial reads in HUVECs infected with UT176 (Fig. 1D, Supp. Fig. 3). This difference likely reflects growth rate differences between Karp and UT176, which have doubling times of 19 and 27 hours in HUVEC, respectively (Fig. 1B). The distribution of reads to RNA classes (Fig. 1D) indicated efficient depletion of ribosomal transcripts in the host transcriptome (<0.001% human rRNA reads). In contrast, we found an average of 16% and 30% rRNA reads in Karp and UT176, respectively. Most of these remaining bacterial ribosomal reads were derived from 5S rRNA (Supp. Table), likely reflecting the divergence of 5S rRNA sequences between Ot and bacterial model organisms used for optimization of the Ribo-Zero approach (https://emea.illumina.com/products/selection-tools/ribo-zero-kit-species-compatibility.html?langsel=/de/). This notwithstanding, reads mapping to coding sequences (CDSs) were not only abundant in the HUVEC data subset (53% of all host-mapped reads), but also in the Ot-specific reads (38% of the Karp- and 49% of the UT176-mapped reads), allowing differential expression analysis. Dual RNA-seq also readily detected the various non-coding RNA classes from both host and bacteria (Fig. 1E). Of 657 predicted core Ot genes^8^ 491 were expressed and 212 were highly expressed (see Methods for definitions).

### Ot ncRNAs and evidence for tmRNA processing

Bacterial genomes encode many non-coding (nc)RNAs. Among the most conserved are several specialized, abundant housekeeping ncRNAs, including the RNA components of ribonuclease P (RNase P), the signal recognition particle (SRP), and transfer-messenger RNA (tmRNA), all of which were detected in the Karp transcriptome data (Fig. 1E, Supp. Table 1). To validate the RNA-seq data, we performed Northern blot analysis for conserved housekeeping ncRNAs (Fig. 2A). These include the M1 RNA component of RNase P, a ribozyme responsible for tRNA processing, and 4.5S, the RNA component of the SRP involved in translocation of membrane proteins. Both ran at their expected lengths of ∼385 and ∼100 nt, respectively. However, a second stronger band for the M1 transcript ran slightly higher, indicative of a length of ∼450 nt, suggesting the existence of a precursor-M1.

**Figure 2.**
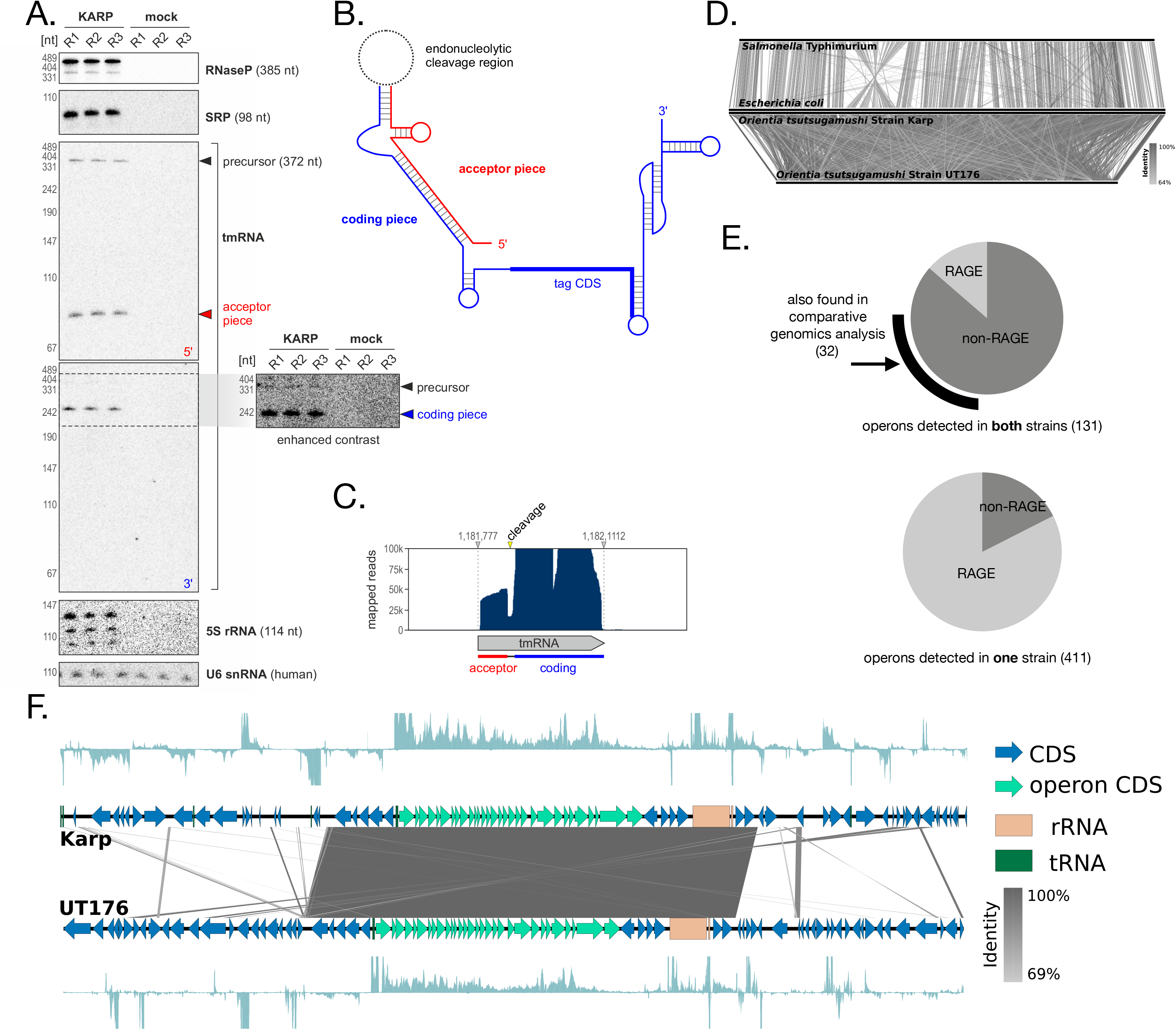
RNA biology in Ot. **A.** Northern blot analysis of core non-coding RNAs in Ot. **B.** Structure of the two-piece tmRNA observed in the Ot transcriptome **C.** RNA-seq read coverage over the tmRNA gene mirrors cleavage observed by Northern blot. **D.** A comparison of genomic synteny of two species within the enterobacteriaceae (*Escherichia coli* MG1655 and *Salmonella enterica* serovar Typhimurium SL1344, top), with synteny between the two Orientia strains from this study (bottom). **E.** Pie charts illustrating the relative abundance of RAGE genes in conserved (top) and strain-specific (bottom) operons. **F.** Visualization of the largest conserved operon in Ot, encoding multiple ribosomal genes, showing RNA-seq coverage in both strains.

We also found evidence of tmRNA processing in Ot. tmRNA has both mRNA-like and tRNA-like features, rescues stalled ribosomes^30^, and is known to contribute to virulence in pathogens as diverse as *Salmonella* Typhimurium^31^ and *Francisella tularensis*^32^. In our data, tmRNA appears to be expressed at unusually high levels, contributing between 3 and 7% of total bacterial reads (Fig 1E), suggesting an important role in Ot survival in mammalian cells. tmRNA generally consists of a tRNA-like (acceptor) domain encoded upstream of a short open reading frame (coding domain). However, the transcript has undergone a circular permutation in some clades of bacteria^33^, including the Alphaproteobacteria^34^, which requires processing of a precursor transcript into separate, base-pairing acceptor and coding RNA chains^35, 36^ (Fig. 2B). We detected three Ot tmRNA forms using Northern blot: (i) a long precursor tmRNA (372 nt); (ii) a 5’ fragment of ∼80 nt, the acceptor domain; (iii) and the 3’ coding domain of ∼240 nt (Fig. 2B). Read coverage over the tmRNA locus in the Karp genome supported a cleavage event within the loop region that connects the tRNA- and mRNA-like domains in the full-length precursor (Fig. 2C).

In addition to these universally conserved housekeeping ncRNAs, bacterial genomes encode family-, genus-, species-, or strain-specific small ncRNAs (sRNAs) to adapt their gene expression to specific intrinsic and environmental cues^37, 38^. Our RNA-seq data identified 55 intergenic sRNA candidates, between 77-803 nt, in the Karp transcriptome (Supp. Table 1). When normalized to the genome size of Ot, this is consistent with the number of sRNAs reported in model bacterial pathogens^39–44^.

### Conserved operons in a dynamic genome

The genome of Ot is highly dynamic^8^, and while the timescales and mechanisms of its rearrangements are unknown they are thought to be driven by an extreme proliferation of mobile elements^6, 7^, in particular the RAGE. The consequences of this are evident when comparing the high degree of synteny in bacteria from two related ‘normal’ genera (*Escherichia* and *Salmonella*) to the complete shuffling we observe between the two Ot strains studied here (Fig. 2D). As bacterial genomes are normally organized into co-transcribed operons of functionally related genes, we wondered how this macroscale loss of synteny would affect conservation of these transcripts. Using Rockhopper^45^ and manual curation, we identified adjacent genes expressed in a continuous transcript, classifying these as operons. We identified 131 operons fully conserved between Karp and UT176 (all genes expressed in both strains) and seven partly conserved (some genes expressed in both strains). Our previous analysis of 8 Ot genomes identified 51 universally conserved genomic islands, including 35 potential collinear gene clusters containing two to thirteen genes^8^, and we found evidence for operonic transcripts originating from 24 of these. We also identified 212 and 192 transcribed operons present only in Karp or UT176, respectively, and these were generally associated with the RAGE mobile element (73% in Karp and 93% in UT176) in contrast to conserved operons (14% of fully conserved operons, Fig 2E).

The majority (84%, Supp. Fig. 6) of conserved operons consisted of only two or three genes. Longer operons tended to encode for core cellular processes, the longest being a 30 gene operon encoding almost half of Ot ribosomal proteins proximal to the ribosomal RNA operon itself (Fig. 2F). Others included an 8 gene operon involved in iron-sulfur cluster assembly, and 6 and 5 gene operons in distinct loci each encoding for portions of the NADH-ubiquinone oxidoreductase complex in an organization similar to that observed in *Rickettsia prowazekii* and eukaryotic mitochondria^46^. In summary, the identification of cotranscribed gene clusters in a genome as highly dynamic as that of Ot indicates strong selection for those genes to remain coupled, indicating involvement in the same pathways and likely shared regulation.

### Evidence for Ot RAGE regulation by antisense RNA

The RAGE of Ot is present in at least 185 remnant copies^7^. It encodes an integrase (*int*) and transposase gene *(tra),* multiple genes from the VirB type IV secretion system (*vir*) and a number of potential effector genes including ankyrin-repeat containing proteins (*ank),* tetratricopeptide repeat containing proteins (*TPR*), SpoT/RelA genes, DNA methyltransferases and replicative DNA helicases. Many of these genes are truncated and most RAGE copies are highly degraded, containing only a subset of genes from the complete element. It is not known if this ICE is still active for transposition, nor whether Ot can express a functional type IV secretion apparatus. In our RNA-seq dataset ∼50% of the most highly expressed genes were repetitive genes encoded by the RAGE (defined throughout our analysis as integrase, transposase, conjugal transfer genes and hypothetical genes) in both strains. These same genes were also highly expressed in the antisense direction (Fig. 3A, Supp. Fig. 6), leading us to hypothesize that the repetitive RAGE genes may be regulated by antisense gene expression.

**Figure 3.**
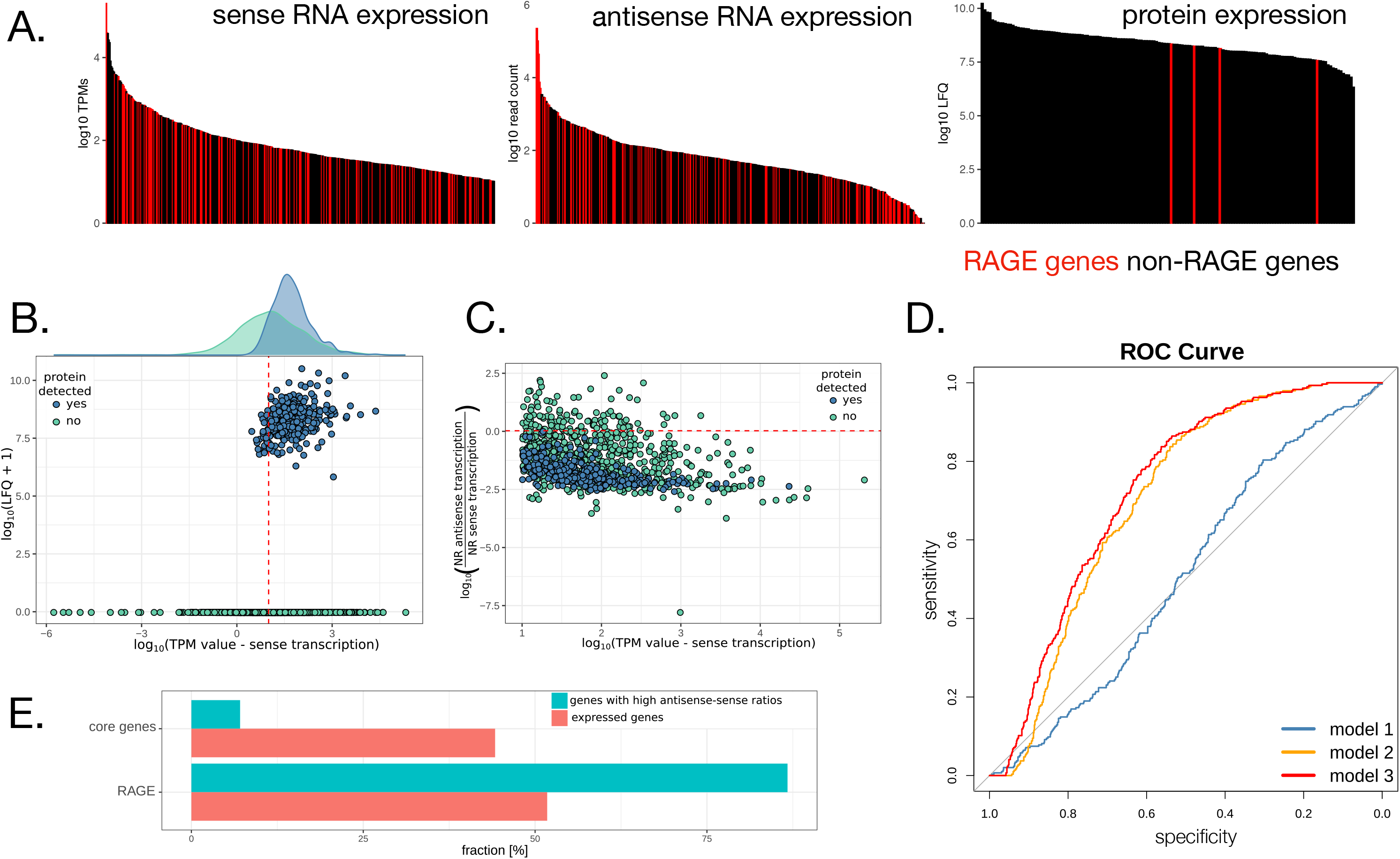
Antisense transcription is enriched on mobile genetic elements and is predictive of an absence of protein expression. **A. B.** Plot showing the relationship between protein expression, defined by LFQs, and transcript expression, defined by TPMs. Genes cluster into two groups based on their protein expression. The red line indicates the threshold for expressed genes (TPM value equal to 10). **C.** Sense transcription and the ratio of reads assigned to the antisense and sense strands, showing classification based on proteomics detection. The red line indicates the sense-antisense ratio (1.059182) above which translation was not detected by mass spectrometry. **D.** ROC curves evaluating the performance of logistic regression models to predict protein expression from RNA-seq read counts. Model 1 strictly uses sense expression, Model 2 the antisense-sense ratio, and Model 3 uses both. Incorporating antisense expression clearly improves model performance. **E.** Fraction of core genes and RAGE genes in the set of genes with high antisense-sense ratios, compared to all expressed genes.

Antisense transcription is widespread in bacteria^47^, with between 5% and 75% of coding sequences exhibiting antisense transcription. While functions for a number of specific antisense transcripts have been described, including regulation through occlusion of the ribosome binding-site or induction of RNase III mediated decay^48^ their relevance as a general functional class remains unclear. Antisense promoters tend to be weakly conserved^49^, arguing against specific functions, and mathematical modeling has suggested the majority of antisense transcripts are not expressed at sufficient levels to affect the regulation of their cognate coding sequence^50^.

To explore the relationship between sense and antisense expression of core Ot genes and repetitive RAGE genes, we combined our Karp RNA-seq dataset with a proteomics dataset generated under the same experimental growth conditions. We observed substantially fewer RAGE gene products detected by proteomics, compared with RNA-seq (Fig. 3A). Genes with detected protein products had higher transcript expression on average compared to those not detected by proteomics (Fig. 3B). However, many highly expressed transcripts appeared to produce no protein. Given our previous observations, we asked whether antisense transcription would correlate with protein expression. All genes with detected proteins had an antisense-sense read count ratio of less than 1, in contrast to genes with no detected protein product, which had an antisense-sense read count spanning several orders of magnitude (Fig. 3C) suggesting antisense RNA expression may be a factor in inhibiting translation.

To test this hypothesis more rigorously, we constructed three logistic regression models to predict protein detection from our transcriptomic data. The first used only transcripts per million (TPMs) derived from the sense strand as a predictor; the second used only the antisense-sense read count ratio as a predictor; the third used both features. Comparisons of the predictive power of these three models showed that antisense transcription is predictive of protein expression (Fig. 3D). Model 1, relying only on sense expression, did little better than chance at predicting protein detection. Models 2 and 3, which incorporate the antisense-sense ratio, led to large improvements in predictive power, suggesting that antisense transcription plays a wide-spread regulatory role in Ot. This was confirmed by cross-validation (Methods, Supp. Fig. 7). We found significant enrichment for RAGE genes among those with high antisense-sense ratios (Fig. 3E), suggesting antisense transcription may work to control the expression of selfish genetic elements at the protein level. Thirty core genes also exhibited an antisense-sense ratio of >1 (Supp. Table core genes) and these include the chromosomal replication initiator protein DnaA, DNA polymerase subunit III, an outer membrane autotransporter protein ScaD, glutamine synthetase, three transporters, the protein export protein SecB and 12 hypothetical proteins. None of these models achieved greater than 67% balanced accuracy, which may be due to both the existence of other modes of post-transcriptional regulation and the lack of sensitivity in our proteomics. For instance, we have also performed a preliminary investigation of codon bias and found some evidence for differential codon usage in genes expressed at the RNA, but not protein, level (Supp. Text, Supp. Fig. 8-9).

### Differential expression of genes in Karp and UT176

Due to a lack of genetic tools, identification of virulence mechanisms in Ot has been difficult, with only a small number of antigenic surface proteins and effectors known. As pangenome diversity appears to primarily be the result of gene duplication and decay, differences in virulence between strains are likely due to differences in expression. To investigate this hypothesis, we performed differential expression analysis between Karp and UT176 at 5 d after infection of HUVEC cells. Pathway and gene ontology (GO) analyses of differentially expressed genes (Fig. 4A, Supp. Table 1) indicated that most pathways were up-regulated in Karp compared with UT176, including those involved in DNA replication and metabolism, consistent with Karp’s higher growth rate (Fig. 1B). At the gene level (Supp. Table 1, Fig. 4B) we found a number of surface and effector proteins (Anks) were differentially regulated between the two strains. Ot encodes five autotransporter domain-containing proteins (ScaA-ScaE) and three immunogenic-type surface antigens (TSA22, TSA47, TSA56). All these surface proteins are immunogenic, based on their reactivity to patient sera^60^, with TSA56 being the most abundant Ot surface protein. TSA56 has four variable domains and these lead to strain-specific antibody responses in patients. TSA47, TSA56, and ScaA have been evaluated as possible vaccine candidates^61, 62^. Of the core Ot genes, those most differentially expressed between Karp and UT176 were *scaE*, *tsa56,* and *tsa22* (1.39, 3.06 and 3.94 logFC in Karp over UT176, respectively). In contrast, *scaD* levels were increased in UT176 but to a lesser degree (0.80 logFC in UT176 over Karp). Given their high immunogenicity it is likely that their differential expression results in differential host responses.

**Figure 4.**
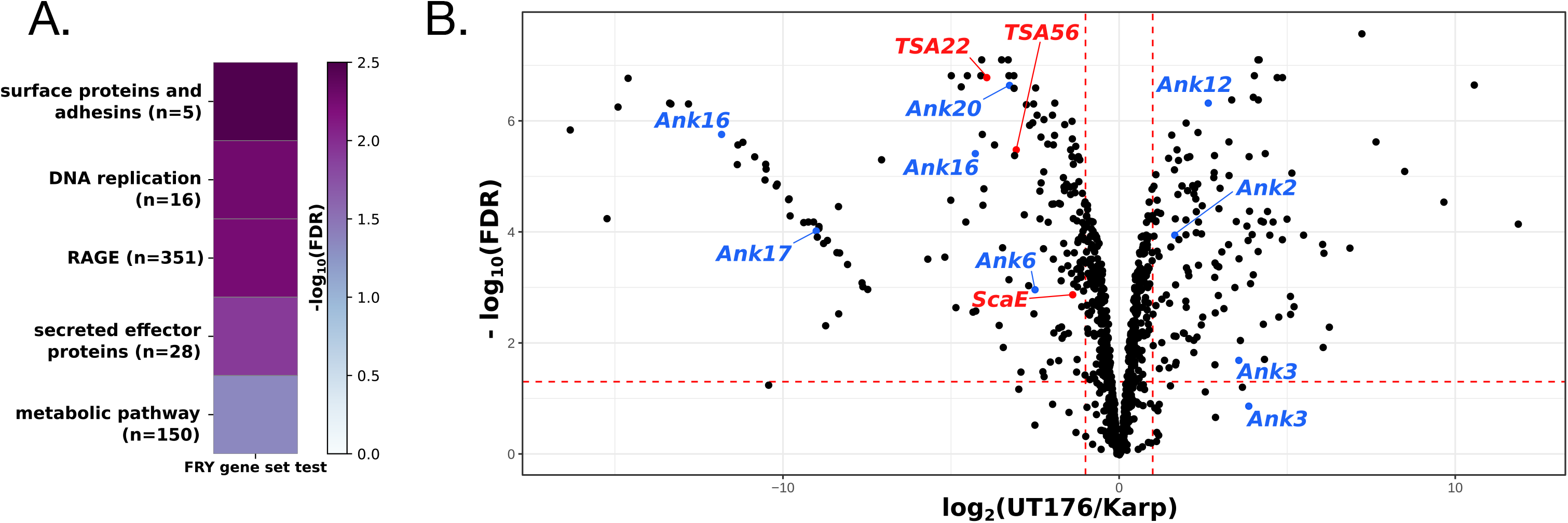
Differential bacterial gene expression. **A.** Heatmap illustrating pathways enriched in differentially expressed genes. All illustrated categories are more highly expressed in Karp. **B.** Volcano plot showing the differential expression of bacterial genes in Karp and UT176. Bacterial surface genes (red) and ankyrin repeat-containing effector proteins (blue) with log fold change ≥1 are highlighted.

Genes for Ank and tetratricopeptide repeat-containing proteins (TPR) are present in 33 (Ank)/29 (TPR) and 21 (Ank)/22 (TPR) copies in Karp and UT176, respectively ^8^. Some *ank* genes function as effectors in eukaryotic cells whilst others are uncharacterized. We compared the expression of Ank and TPR genes in Karp and UT176. *ank2*, *ank3*, *ank12*, *tpr6,* and *tpr8* were up-regulated with a logFC >1.5 in UT176, while *ank16*, *ank17*, *ak20,* and *tpr1* were up-regulated with a logFC >1.5 in Karp. Most of these proteins were not detected in the Karp proteomics dataset suggesting that either the mRNAs were not translated, or that the proteins were secreted and lost during purification. The protein products of all of these *ank* genes localise to the endoplasmic reticulum or host cell cytoplasm when ectopically expressed^63^. Ank6 interferes with NFkB translocation to the nucleus and inhibits its transcriptional activation^64^. The activity of the other differentially expressed Anks is not known. Given that these effector proteins interact directly with host cell proteins we expect that this differential expression will lead to downstream differences in host response.

### Karp and UT176 induce a type-I interferon proinflammatory response

The transcriptional profile of HUVEC cells infected with Karp or UT176 showed a clear core response to Ot (Fig. 5A, red), with smaller gene sets responding specifically to a single strain (Fig. 5A purple and orange). The core response was dominated by a type-I interferon proinflammatory response (Supp. Table 1), seen previously in cultured endothelial cells and monocytes, as well as patient-derived macrophages^13–17^. This is further illustrated by activation of the canonical interferon signaling pathway in response to Karp (Fig. 5B), with a similar response observed for UT176 (Fig. 5C).

**Figure 5.**
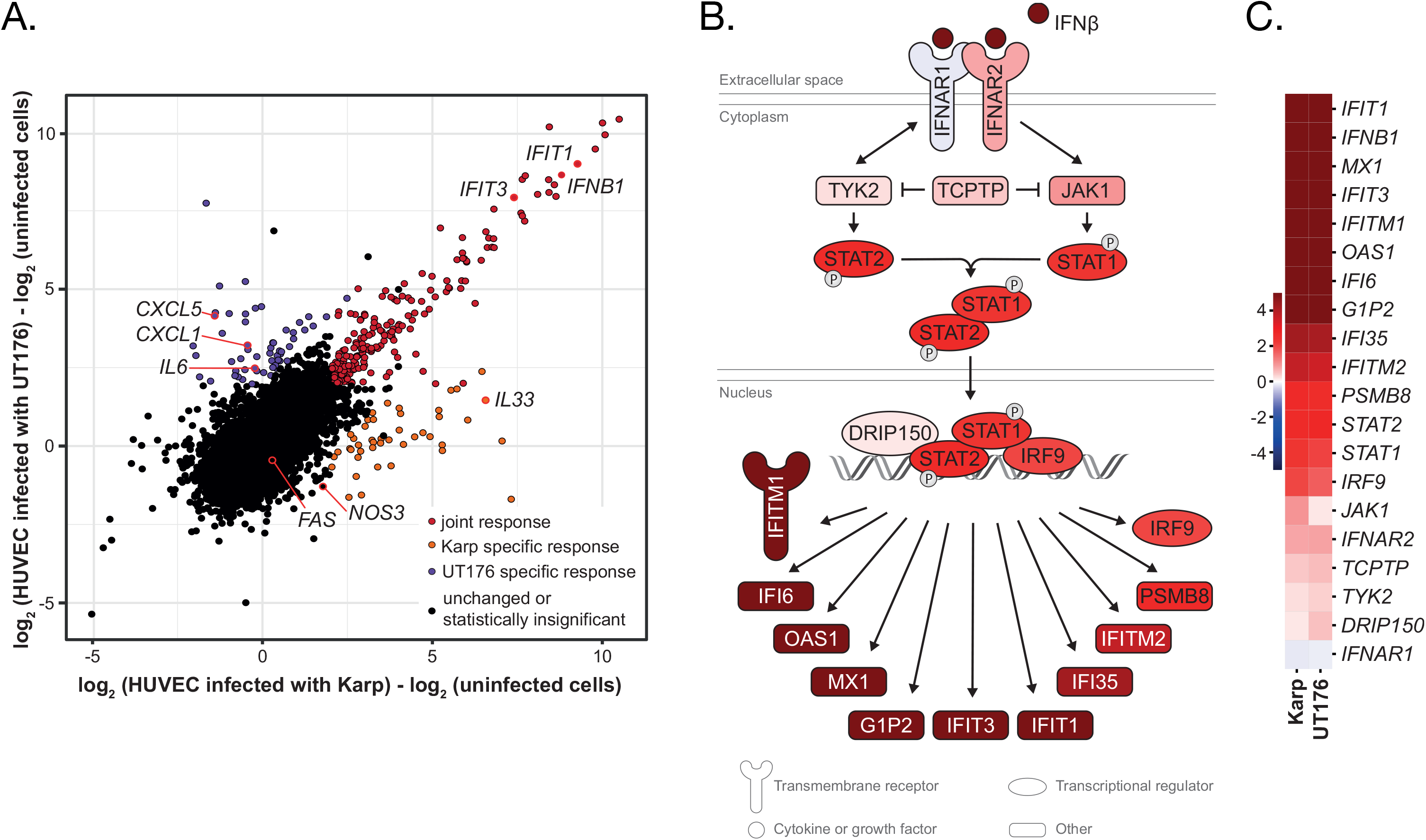
Ot induces an antiviral interferon response in HUVECs. **A.** Summary of the host response showing joint and strain specific responses. The joint response is defined as genes with a logFC > 2 and FDR-corrected p-value < 0.01 for infection with both Karp and UT176. Strain-specific responses are genes with a logFC > 2 and FDR-corrected p-value < 0.01 for infection with either Karp or UT176, excluding genes already included in the joint response. **B.** Activation of multiple genes in the canonical interferon signaling pathway in Karp infected HUVECs compared with uninfected HUVEC cells. **C.** Heat map showing up-regulation of genes in the interferon signaling pathway in HUVEC cells infected with Karp and UT176 compared with uninfected cells.

Host genes commonly up-regulated upon infection with either Ot strain include *IFNB1* (interferon beta) and genes involved in regulating the type-I interferon response: *IRF9* (interferon-regulated factor 9) and *STAT1*/*2*. Interferon-stimulated genes were also up-regulated upon Ot infection, including various interferon induced proteins with tetratricopeptide repeats (*IFIT*) genes and 2’-5’-oligoadenylate synthase 1 (*OAS1*). In addition to the type-I interferon pathway, the joint Ot response led to up-regulation of proinflammatory chemokine genes including *CXCL10*, *CXCL11*, and for cytokine receptors *IL13RA2*, *IL7R*, *IL15RA*, and *IL3RA* (Supp. Table 1).

The upstream signals leading to activation of these signaling pathways are unknown but Ot has been shown to activate host cells by signaling through the NOD1-IL32^65^ and TLR2^66^ pathways. Our data showed that *TLR3* is up-regulated in cultured HUVEC cells in response to both Karp and UT176 (Supp. Table 1). TLR3 recognizes viral double-stranded (ds)RNA in the cytoplasm^67^ and it is possible that it responds to Ot dsRNA. The up-regulation of the mRNA for transcription factor IRF7, which is known to respond to stimulation from membrane-bound TLRs, further supports a role for TLR2 and TLR3 in the detection of Ot.

### Differential host responses to Karp and UT176

Although Karp and UT176 both induced a type-I interferon proinflammatory response compared to uninfected HUVEC cells (Fig. 5B,C), each strain also induced its own unique response. Some of these expression changes were validated by qRT-PCR (Supp. Fig. 12). The mRNA levels of multiple cytokines, chemokines, and cytokine receptors were higher in HUVEC cells infected with UT176 compared with Karp (Fig. 6A, Supp Fig 14, 15, 17). A network map of proinflammatory chemokines and cytokines, and their differential induction in response to UT176 and Karp is shown in Fig 6A and Supp. Fig. 14A. Most of the genes for cytokines, chemokines, and cytokine receptors were differentially up-regulated by infection with UT176 compared with Karp, including *CXCL8*, *CXCL1*, *CXCL2*, *CXCL10*, *IL6*, *IL1RL1*, *IL18R1*. The mRNA levels of surface adhesion molecules associated with activation of the endothelium, VCAM1 and ICAM1, were also up-regulated in UT176 infected cells compared with Karp (Supp. Table 1). Whilst *TLR3* was up-regulated in both strains, TLR3 activation in UT176 infected cells was 1.5 logFC higher than in response to Karp. Comparison of NFkB pathway genes and genes associated with NOS2 production revealed that genes in both pathways were up-regulated in UT176 but they were not up-regulated or were significantly less up-regulated in response to Karp infection (Supp. Fig. 15). Expression of host genes associated with leukocyte proliferation and mononuclear leukocyte differentiation was strongly induced in HUVECs infected with UT176 but significantly less so when infected with Karp (Supp. Fig. 16). Thus, UT176 seems to induce a stronger proinflammatory response and this may lead to more effective pathogen clearance (see Fig. 1B).

**Figure 6.**
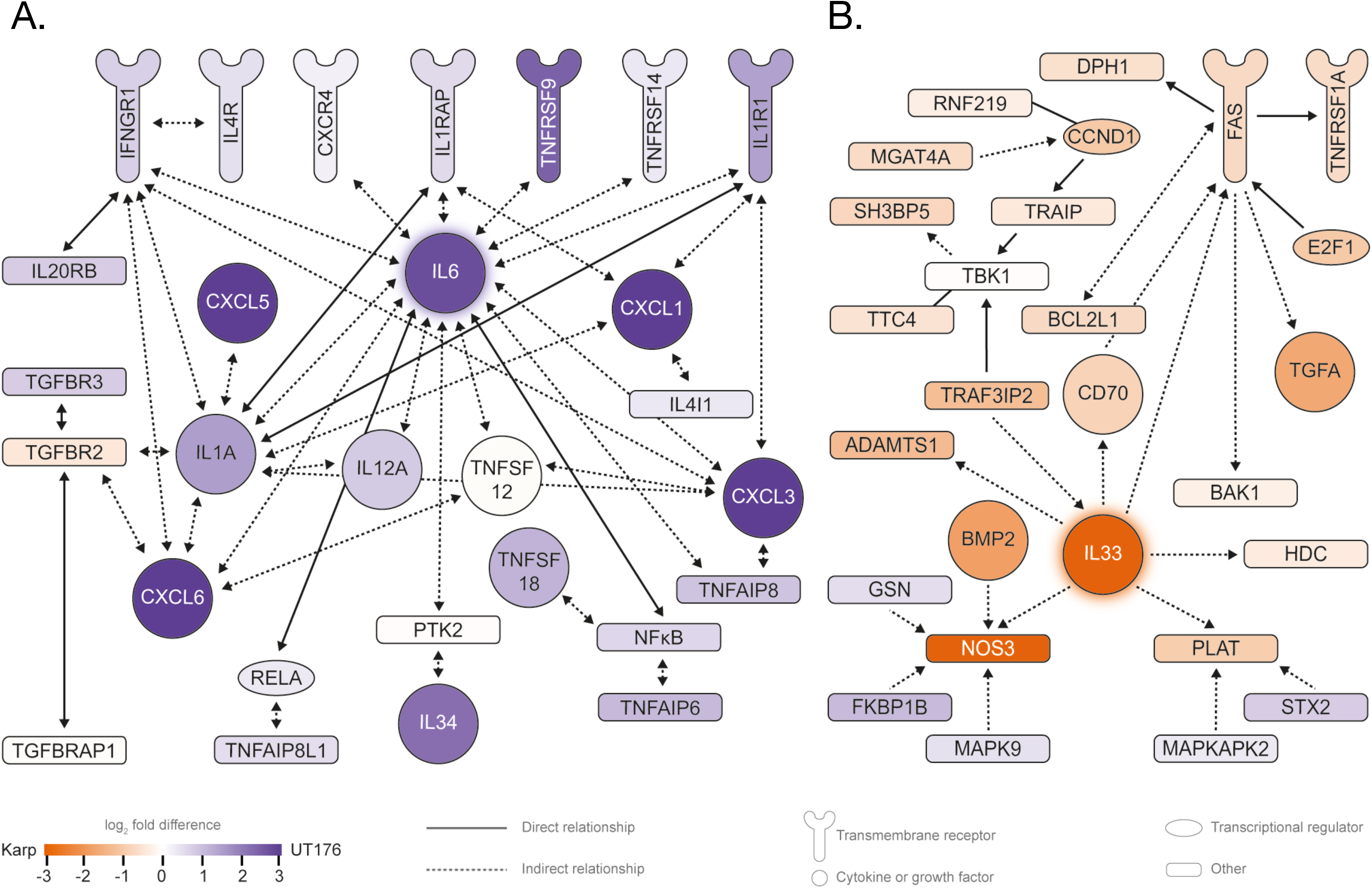
Karp and UT176 lead to the up-regulation of distinct networks in HUVECs. **A.** Up-regulation of multiple proinflammatory chemokines and cytokines in HUVECs infected with UT176. **B.** Induction of the IL33-FAS-mediated anoikis network in Karp infected HUVECs.

In contrast to the multiple chemokines and cytokines up-regulated in UT176-infected HUVEC cells, only *IL33* was specifically up-regulated in Karp-infected HUVEC cells (5 logFC difference; Supp. Table 1). IL33 is a proinflammatory cytokine that is involved in pathogenicity in a mouse model of scrub typhus^68^. To investigate Karp-mediated activation of *IL33* we analyzed gene induction in the IL33-FAS network (Fig. 6B/Supp. Fig. 14B). Most genes in the network were differentially induced in Karp-infected HUVEC cells compared to in UT176-infected HUVECs. Up-regulation of IL33-NOS-mediated signaling contributes to tissue inflammation. We analyzed networks of genes involved in (i) organismal growth failure (ii) organismal morbidity and mortality and (iii) organismal death. In all cases Karp induced these networks whilst UT176 dampened them (Supp. Fig. 17).

### Two Ot strains differ in virulence in a mouse model

To assay the effects of the differential inflammatory responses induced by Karp and UT176 we tested the relative virulence of the two strains in an intravenous mouse infection model. 1.25×10^6^ bacteria were inoculated into outbred ICR mice and monitored for disease symptoms for 12 days prior to euthanasia. Blood and tissue from lung, liver, spleen, and kidneys were isolated and the bacterial load measured by qPCR. We found that Karp was significantly more virulent than UT176 as determined by disease outcome, weight loss, bacterial load, and histopathological analysis (Fig. 7, Supp. Fig. 19-21). This difference is likely to result from a combination of differential bacterial growth rate (Fig. 1B), differential expression of bacterial virulence related genes (Fig. 4), and differential induction of host immune networks (Fig. 6).

**Figure 7.**
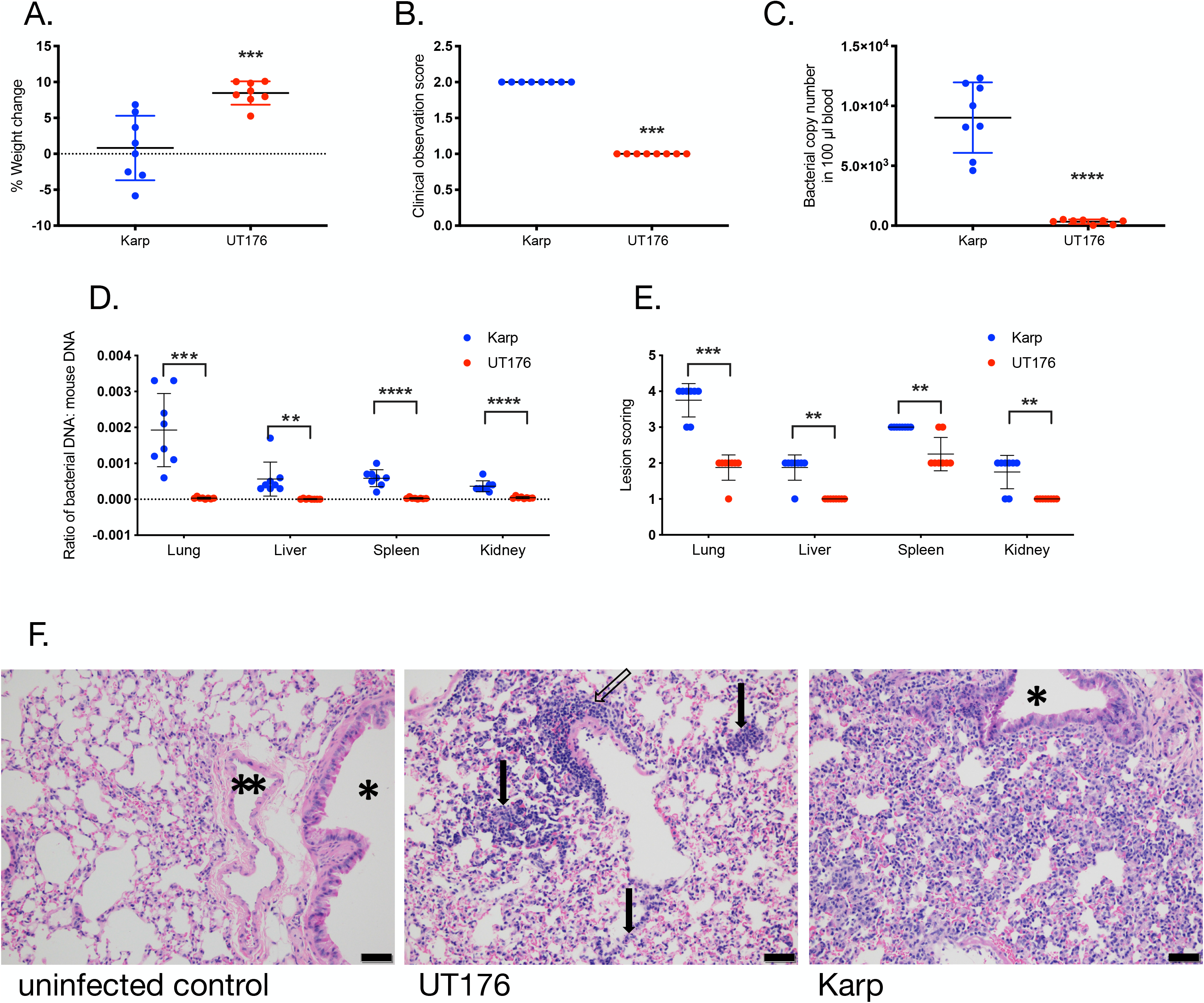
Karp is more virulent than UT176 in a mouse infection model. **A.** Weight change over 12 days of infection. **B.** Clinical observation score of mice 12 days post-infection. This number is a composite score based on appetite, activity, and hair coat with higher numbers representing low appetite, low activity, and ruffled fur. Details provided in Supp. Fig. 20. **C.** Bacterial copy number in 100 µl blood taken from euthanized mice 12 days post-infection, measured by qPCR. **D.** The ratio of bacterial DNA copy number to mouse DNA copy number in lung, liver, spleen, and kidney of euthanized mice 12 days post-infection, measured by qPCR. **E.** Lesion scores of Hematoxylin and Eosin stained lung, liver, spleen, and kidneys of euthanized mice 12 days post infection. Scores range from 0 to 5 with 0 representing normal tissue and 5 representing severe lesion damage. All graphs show mean and standard deviation. Statistical significance is calculated using unpaired Student t-test in GraphPad Prism software. **p≤0.01 ***p≤0.001 ****p≤0.0001 **F.** Images of Hematoxylin and Eosin stained lung tissue of mice infected with buffer, UT176 or Karp. Scale bars = 50 µm. * indicates airway and ** indicates blood vessel. Uninfected control: airway, blood vessel and alveoli all appear normal. UT176-infected lungs: There are diffuse thickening and infiltration of alveolar septa with a mixed population of macrophages and lymphocytes (arrows). There is also mild perivascular lymphohistiocytic inflammation (open arrow). Karp-infected lungs: There is diffuse moderate thickening and infiltration of alveolar septa with a mixed population of macrophages and lymphocytes. The airway (*) is unaffected and normal. Additional figures are shown in Supp. Fig. 20.

## Discussion

Both its obligate intracellular lifestyle and the complexity of rearrangements in the Ot genome make it difficult to study. Ot has a genome of 1.9-2.5 Mbp, almost half of which is composed of repetitive regions of >1,000 bp in length^8^. This is in contrast to the most closely related rickettsial species, whose genomes are typically around 1.1-1.3 Mbp^69^. The Ot genome is remarkably unstable, which makes inference of its transcriptional architecture particularly difficult. Using RNA-seq, we were able to identify core ncRNAs, putative sRNAs, and operonic transcripts. In sharp contrast to most bacteria, only a handful of operons containing more than 2 or 3 genes were conserved between Karp and UT176, and these primarily encode for proteins involved in core cellular processes like respiration and translation. Given that Karp encodes only 12 predicted transcription factors and 3 sigma factors, in contrast to 300 and 7, respectively, in *E. coli*, this raises the question of how transcription in Ot is coordinated.

One possible explanation is that much Ot transcription is not stringently controlled, and alternative mechanisms have arisen in Ot to control protein expression. This is supported in part by our observation that protein expression is partially predicted by antisense transcription. This mode of regulation seems to be particularly prevalent for genes encoded by the RAGE, a transposable element of the integrative and conjugative element group. Transposable element regulation by antisense transcripts was one of the earliest discovered examples of riboregulation^71^, though it has not previously been observed at the scale implied by our RNA-seq analysis. Such antisense regulation could arise spontaneously through capture of transcriptional noise, providing a parsimonious alternative to transcriptional control^72^. It is unclear whether these untranslated transcripts have some function in Ot, or whether they are purely selfish DNA elements that Ot has been unable to dispose of due to its small population size. One intriguing possibility is that this regulatory mechanism would provide a large pool of double stranded RNA upon intracellular bacterial lysis, which may explain Ot induction of TLR3 and an antiviral immune response.

In the absence of genetic tools it is difficult to identify specific genes that drive virulence differences between UT176 and Karp. However, comparative genomics has revealed that whilst the pan genome of Ot is open, it is largely composed of gene duplications rather than newly acquired genes. This lack of gene novelty likely reflects the environmental isolation associated with an obligate intracellular lifestyle. Consequently, strain-specific differences in virulence are likely to be driven largely by differences in relative gene expression rather than the presence or absence of virulence genes. Consistent with this, we observed an up-regulation of virulence-associated surface proteins in Karp compared with UT176.

The inflammatory response triggered by Ot infection is a key driver of virulence in scrub typhus. We compared the response of endothelial cells to the two strains of Ot and found that differential activation of the immune response correlated with differential outcomes in a scrub typhus mouse model. Whilst both Karp and UT176 induced an antiviral proinflammatory response, as shown previously^13–17^, UT176 strongly induced an IL6-mediated pro-inflammatory response whilst Karp induced an IL33-NOS3-FAS response, differences likely to influence the relative virulence of these strains.

*IL33* was one of the most strongly differentially regulated genes between UT176 and Karp infections (5.1 logFC higher in Karp-infected HUVECs). IL33 has previously been shown to play a role in pathogenesis in a scrub typhus murine model, using the Karp strain, where it was shown that IL33 levels were increased during Ot infection, that *IL33*-/- mice showed less severe disease symptoms, and that addition of rIL33 increased severity and mortality^68^. Our observations of reduced induction of *IL33* by the less virulent UT176 strain further support a role for this cytokine in the pathogenesis of scrub typhus.

In summary, we have used dual RNA-seq to gain insights into the transcriptome structure and mechanisms of gene regulation in the neglected intracellular pathogen Ot during infection. We provide evidence for widespread antisense regulation, in particular for the RAGE genes. We identified a relationship between the relative induction of IL33- and IL6-based gene networks in the host and disease severity. These findings will lay the groundwork for subsequent studies on the regulation of gene expression in Ot and mechanisms of pathogenesis. More generally, the present study may serve as a blueprint for the characterization of further obligate intracellular, genetically intractable bacterial pathogens.

## Materials and Methods

### Growth of Ot and isolation of RNA

The clinical isolate strains (Karp and UT176) of *Orientia tsutsugamushi* were propagated in a confluent monolayer of host cells (HUVEC, Human Umbilical Vein Endothelial Cells; ATCC PCS-100-010) for 5 days at MOI 100:1. Cells were cultured using Media200 (Thermo Fisher, Catalog no. M200-500) supplemented with LVES media (Thermo Fisher, Catalog no. A14608-01) at 35 °C and 5% CO_2_. The infectivity was determined by qPCR of the single copy Ot gene *47kDa* at day 5-7. Both uninfected cells and infected cells were harvested by incubating the cells on ice and quickly resuspending in RNAprotect Bacteria Reagent (Qiagen, catalog no. 76506), then storing at −80 °C until use. RNA extraction was performed using the Qiagen RNeasy Plus kit (Qiagen, catalog number 74136) according to manufacturer’s instructions and as described previously (Atwal, 2017).

### RNA processing and sequencing

The integrity of the DNase-treated RNA samples was assessed in a Bioanalyzer (Agilent). All samples had RIN (RNA integrity number) values ≥8.0. Ribosomal transcripts were removed using the Ribo-Zero Gold (epidemiology) kit (Illumina). Following the manufacturer’s instructions, 500 ng of total, DNase-treated RNA was used as an input to the ribo-depletion procedure. rRNA-depleted RNA was precipitated in ethanol for 3 h at −20°C.

cDNA libraries for Illumina sequencing were generated by Vertis Biotechnologie AG, Freising-Weihenstephan, Germany. rRNA-free RNA samples were first sheared via ultrasound sonication (four 30-s pulses at 4°C) to generate on average 200- to 400-nt fragments. Fragments of 20 nt were removed using the Agencourt RNAClean XP kit (Beckman Coulter Genomics) and the Illumina TruSeq adapter was ligated to the 3’ ends of the remaining fragments. First-strand cDNA synthesis was performed using M-MLV reverse transcriptase (NEB) wherein the 3’ adapter served as a primer. The first-strand cDNA was purified, and the 5’ Illumina TruSeq sequencing adapter was ligated to the 3’ end of the antisense cDNA. The resulting cDNA was PCR-amplified to about 10 to 20 ng/µl using a high fidelity DNA polymerase. The TruSeq barcode sequences were part of the 5’ and 3’ TruSeq sequencing adapters. The cDNA library was purified using the Agencourt AMPure XP kit (Beckman Coulter Genomics) and analyzed by capillary electrophoresis (Shimadzu MultiNA microchip).

For sequencing, cDNA libraries were pooled in approximately equimolar amounts. The cDNA pool was size fractionated in the size range of 200 to 600 bp using a differential cleanup with the Agencourt AMPure kit (Beckman Coulter Genomics). Aliquots of the cDNA pools were analyzed by capillary electrophoresis (Shimadzu MultiNA microchip). Sequencing was performed on a NextSeq 500 platform (Illumina) at Vertis Biotechnologie AG, Freising-Weihenstephan, Germany (single-end mode; 75 cycles).

### Northern blots

Each 15 µg of total RNA (i.e. a mixture of human and Ot RNA) prepared as above were loaded per lane and separated in 6% (vol/vol) polyacrylamide–7 M urea gels. Blotting was performed as previously described^24^. After the transfer onto Hybond XL membranes (Amersham), RNA was cross-linked with UV light and hybridized at 42°C with gene-specific 32P-end-labeled DNA oligonucleotides (Supp. Fig. 13) in Hybri-Quick buffer (Carl Roth AG). After exposure, the screens were read out on a Typhoon FLA 7000 phosphorimager (GE Healthcare).

### qRT-PCR

qRT-PCR was performed with the Power SYBR Green RNA-to-CT1-Step kit (Applied Biosystems) according to the manufacturer’s instructions and a CFX96 Touch real-time PCR detection system (Bio-Rad). Human U6 snRNA served as reference transcripts. Fold changes in expression were determined using the 2(–ΔΔCt) method ^76^. Primer sequences are given in Supp. Fig. 13, and their specificity had been confirmed using Primer-BLAST (NCBI).

### RNA seq read processing and quantification

The raw reads were initially processed according to our established dual RNA-seq pipeline^24^. Briefly, raw reads were trimmed for adaptor sequences and a minimum read quality of 20 using cutadapt^77^. Reads were then mapped against the human (GRCh38) and Ot (UT176 accession: LS398547.1; Karp accession: LS398548.1) reference sequences using the READemption pipeline (v0.4.3, ^78^) and segemehl with the lack remapper (v0.2.0 ^79^), removing reads that mapped equally well to the bacterial and host genomes. For downstream analysis of human gene expression, only uniquely mapping reads were retained for quantification.

To improve quantification of repetitive sequences, reads mapped to the Ot genomes were used for quantification of bacterial transcript expression using Salmon (v0.9.1) ^29^. Salmon is a quasi-mapping based gene expression quantification tool that consists of two steps, indexing and quantification. Transcript fasta files were created from the Genbank annotations using the gene coordinates. The indexing step was performed in quasi-mapping mode (--type quasi). Expression of the transcripts was quantified using both stranded forward library type (-lSF) and removing incompatible mappings (--incompatPrior 0.0). Salmon identified identical gene repeats that are collected in 218 groups (see supplementary table, Karp groups of duplicates). For quantification purposes we retained a single gene from each group. For the purposes of summarizing gene expression, we calculated mean TPM values from three replicates for each strain. Genes with a mean TPM greater than 10 were classified as expressed, and those with a mean TPM value greater than 50 highly expressed.

### Gene annotation

For each gene we retrieved the gene name, gene product, and amino acid sequence from the Genbank annotation. In addition, using eggNOG-mapper^10^ we predicted gene names and both KEGG pathways ^11^ and GO terms. We manually identified surface antigen encoding proteins using BLAST. The KEGGREST (Tenenbaum, D (2019) KEGGREST: Client-side REST access to KEGG. R package version 1.18.1.) and GO.db (Carlson M (2019). GO.db: A set of annotation maps describing the entire Gene Ontology. R package version 3.5.0.) R packages were used to retrieve KEGG and GO terms, respectively.

### Non-coding RNA prediction

Noncoding RNAs were annotated using Rockhopper^76^, ANNOgesic ^77^ (v0.7.17) and Infernal^78^ (v1.1.2) searching sequences against the Rfam database^79^. These provided inconsistent predictions of intergenic sRNAs. Intergenic sRNAs were manually curated by visual comparison of the predicted sRNA coordinates with the read coverage in the Integrative Genomics Viewer ^80^ (v2.5.2). Infernal predicted the core housekeeping ncRNAs tmRNA, RNaseP, SRP and 5S rRNA. The quantification of the bacterial transcriptomes complemented with predicted sRNAs was performed using Salmon.

### Genomic alignment

Genomic comparisons in Figure panels D and F were performed using Easyfig^81^. Escherichia coli K-12 MG1655 (Accession number U00096) and Salmonella enterica serovar Typhimurium SL1344 (Accession number FQ312003) were used as comparators for synteny analysis.

### Orthology and conserved operon prediction

We predicted orthologous genes between the two Orientia strains using Poff^82^ with default parameters in synteny mode. To identify conserved operons we used operon structures predicted in each strain by Rockhopper ^41^. Based on visual analysis of read coverage in the Integrative Genomics Viewer, some of the operons were manually extended by addition of genes or merging two operons into one. We also identified partially conserved operons missing some genes in one strain.

### Differential gene expression

For the bacteria, differential gene expression analysis was performed between orthologous genes identified by Poff. Genes that were predicted as an orthologous group (more than two genes) were removed from the analysis. Additionally, we removed duplicates (transcripts with perfectly identical sequence) that were identified by Salmon in either strain. For both human and bacterial RNA-seq data, we performed differential gene expression analysis with the edgeR package ^83^ (v3.20.9) using robust quasi-likelihood estimation ^84^, including genes with CPM (Counts Per Million) > 10 (for Ot) or CPM > 1 (for HUVEC) in at least 3 libraries. To identify biological processes that differ between two Orientia strains, we have performed gene set analysis using KEGG and GO terms that contain at least 4 expressed genes using the fry test in the edgeR package.

### Proteomic sample preparation

Bacteria were propagated in HUVEC cell line at MOI 100:1 and harvested at 5 dpi. Ot was isolated, washed with 0.3 M sucrose, and lysed with 1% triton-X prior to acetone precipitation of protein. Total protein was then alkylated, reduced, and subsequently treated with Lys-C/Trypsin. Digested peptides were desalted using Oasis® HLB reversed-phase cartridges, vacuum dried, and stored for MS runs.

### Mass spectrometry

The dried samples were resuspended of 2% (v/v) acetonitrile solution containing 0.06% (v/v) trifluoroacetic acid and 0.5% (v/v) acetic acid and loaded onto an autosampler plate. Online chromatography was performed using EASY-nLC 1000 (Thermo Scientific) in single-column setup using 0.1% formic acid in water and 0.1% formic acid in acetonitrile as mobile phases. using reversed-phase C18 column (EASY-Spray LC Column, 75 µm inner diameter x 50 cm, 2 µm particle size) (Thermo Scientific). The samples were injected and separated on the analytical column maintained at 50 °C using a 2–23% (v/v) acetonitrile gradient over 60 min, then ramped to 50% over the next 20 min, and finally to 90% within 5 min. The final mixture was maintained for 5 min to elute all remaining peptides. Total run duration for each sample was 90 min at a constant flow rate of 300 nl/min.

Data was acquired using an Orbitrap Fusion mass spectrometer (Thermo Scientific) in data-dependent mode. Samples were ionized 2.5 kV and 300 °C at the nanospray source and positively-charged precursor MS1 signals were detected using an Orbitrap analyzer set to 60,000 resolution, automatic gain control (AGC) target of 400,000 ions, and maximum injection time (IT) of 50 ms. Precursors with charges 2–7 and having the highest ion counts in each MS1 scan were further fragmented using collision-induced dissociation (CID) at 35% normalized collision energy and their MS2 signals were analyzed by ion trap at an AGC of 10,000 and maximum IT of 35 ms. Precursors used for MS2 scans were excluded for 90 s in order to avoid re-sampling of high abundance peptides. The MS1-MS2 cycles were repeated every 3 s until completion of the run.

Identification of proteins within each sample was performed using MaxQuant (v1.5.5.1). Raw mass spectra were searched against Orientia tsutsugamushi primary protein sequences derived from complete genome data. Human whole proteome sequences were obtained from Uniprot and included as background. Carbamidomethylation on Cys was set as the fixed modification and acetylation on protein N-terminus and oxidation of Met were set as dynamic modifications for the search. Trypsin was set as the digestion enzyme and was allowed up to 3 missed cleavage sites. Precursors and fragments were accepted if they had a mass error within 20 ppm. Peptides were matched to spectra at a false discovery rate (FDR) of 1% against the decoy database.

### Proteomic data analysis

Protein expression was measured by label free quantification values (LFQs). A protein was classified as detected if at least two peptides were detected in at least 2 biological replicates, and the mean LFQ across the three replicates was used for further analysis. Otherwise the protein was classified as undetected, and the LFQ value was set to zero. The proteomic data includes 19 protein groups that couldn’t be resolved, consisting of 93 proteins. In our analysis we discarded these proteins to simplify the analysis.

### Transcript classification

Sense transcript expression was defined by mean TPM value across replicates. The antisense/sense ratio was calculated as the ratio of mean read counts assigned to the antisense and sense strand of coding annotations. The duplicated sequences identified by Salmon (Supp. Table 1. Karp groups of duplicates) and non-coding RNAs were removed from the analysis.

We divided the data set into two classes, detected and undetected in proteomics. Within our analyzed data set, 322 genes were detected, whereas 1608 genes were not detected by mass spectrometry. We found a weak positive correlation between TPMs and LFQs for genes with detected proteins (Spearman’s correlation coefficient equal to 0.32), but it was not a linear association (Pearson’s correlation coefficient equal to 0.04). For the further analysis we selected transcripts with sense expression greater than 10 TPMs, previously defined as our expression threshold.

### Logistic Regression model

To test whether antisense-sense ratios are predictive of protein expression, we have applied logistic regression, which models the probability of a binary response, that is, whether a protein is expressed or not. We have built 3 competing models. Model 1 makes predictions of the protein expression based solely on sense transcription:

β_0_ + β_1_* (TPM sense)

Model 2 makes predictions solely on the antisense-sense ratio:

β_0_ + β_1_* ((number of antisense reads) / (number of sense reads))

Model 3 uses both sense transcription and the antisense-sense ratio to make predictions:

β_0_ + β_1_* (TPM sense) + β_2_* ((number of antisense reads) / (number of sense reads))

Since the data is highly imbalanced, 295 transcripts with detected proteins, and 814 without, we used a downsampling procedure (downSample function) implemented in the caret R package^85^ to create a balanced data set for model training purposes. Next, the function glm() with a logit link function from the caret package was used to fit models to the reduced data set. For a first indication as to whether any of these models are predictive, we trained all three models on a downsampled data set consisting of 590 genes, then tested them on the complete data set. To more rigorously assess this result, we have applied 500-fold cross validation. For each fold the data was split randomly into 2 data sets, training and testing which included 1055 and 54 genes, respectively. Each time the new training data set was reduced to 562 genes, which were used to estimate the model parameters, and then the model was evaluated on the testing data set. The model performance was evaluated using a variety of measures, i.e. sensitivity, specificity, accuracy (caret R package) as well as with ROC curves ^86^ (pROC 1.14.0) and the area under the ROC curve (AUC).

### Immunofluorescence microscopy

The protocol for L-Homopropargylglycine (HPG) incorporation, click chemistry and fluorescence detection were based on recommendations from Click-iT® HPG Alexa Fluor® Protein Synthesis Assay Kits (Molecular probe by Life Technologies). HUVECs were grown on chambered coverslip slides (Ibidi, USA), for 2 days before infection with bacteria at MOI 100:1. To incorporate HPG at times indicated, medium was removed and replaced with L-methionine-free medium (Dulbecco’s Modified Eagle Medium, DMEM, Cat. no. 21013) containing 25 µM HPG for 30 min at 37 °C. Labeled bacteria were washed twice in 1X PBS+ 1mg/ml BSA, pH 7.4 before fixing with 4% formaldehyde and subsequently being permeabilized with 0.5% TritonX for 20 min on ice. After washing with PBS + 1 mg/ml BSA, the Click-iT® reaction cocktail (Click-iT® HPG Alexa Fluor® Protein Synthesis Assay Kits cat. C10428) was incubated with cells for 30 min at room temperature in the dark. The Azide dye (Alexa Fluor®488) was used at a final concentration of 5 µM. After the click reaction, cells were labeled with the actin probe Alexa Fluor® 594 phalloidin at a dilution of 1:40 and the nuclear stain Hoechst diluted to 1:1000 for 30 min at 37 °C. Cells were washed 3X with PBS which was replaced with mounting media after the final wash. Imaging was performed using a Zeiss LSM 7000 equipped with a 63 × 1.4 NA objective lens (Carl Zeiss, USA) and also a Leica SP8 laser scanning confocal microscope.

### Analysis of codon bias

We calculated the RSCU (relative synonymous codon usage) for each codon to quantify genome-wide or gene-specific codon usage bias following^52^. To determine the genomic codon counts for each species and gene set, we parsed nucleotide sequence data and annotation in the GenBank file format, downloaded from the NCBI database. We also obtained tRNA gene copy numbers from the GtRNAdb database^87, 88^, and integrated protein abundance for E. coli K-12 MG1655 data from PaxDB 89.

### Host Network/Pathway analysis

To identify pathways that are affected in Karp and/or UT176 infected host cells, genes differentially expressed with an adjusted p-value of < 0.05 were analyzed using Ingenuity Pathway Analysis (IPA) software (Ingenuity® Systems, Inc. Redwood City, CA)^81^ as described previously^82^. Selected pathways were chosen based on enrichment p-values and activation Z-scores, and served as the basis for Figs 5, 6, and Supp. Fig. 11, 14, 15, 16, and 17.

### Mice and Ethics statement

All animal research was performed strictly under approved Institutional Animal Care and Use Committee (IACUC) protocol by the IACUC and Biosafety Review Committee at the Armed Forces Research Institute of Medical Sciences (AFRIMS) Bangkok, Thailand, an AAALAC International-accredited facility. The protocol number was PN16-05. The animal research was conducted in compliance with Thai laws, the Animal Welfare Act, and all applicable U.S. Department of Agriculture, Office of Laboratory Animal Welfare and U.S. Department of Defense guidelines. All animal research adhered to the Guide for the Care and Use of Laboratory Animals, NRC Publication (8^th^ Edition).

C57BL/6NJcl mice were purchased from Nomura Siam International, Bangkok, Thailand. Mice were housed in an animal biosafety level 2 facility and moved to an animal biosafety level 3 containment 2 days before the inoculation. Mice at 6-8 weeks of age were used in these experiments. Two group of mice (n =8 per group) were intravenously injected in the tail vein with 1.25×10^6^ genome copies of *O. tsutsugamushi* of either Karp strain or UT176 strain. The *O. tsutsugamushi* inoculum was derived from *O. tsutsugamushi*-infected L929 cells. Clinical signs and body weight were evaluated daily. After 12 days post inoculation, all mice were euthanized. Blood and tissue samples including lungs, liver, spleen, and kidneys were collected for bacteria quantification and histopathology.

## Acknowledgements

JS is funded by a Royal Society Dorothy Hodgkin Research Fellowship (DH140154) and this project was additionally funded by a grant from the University of Oxford Medical Sciences Division Medical Research Fund. The authors are grateful to Guy Riddihough from Life Science Editors for editorial support on this manuscript. The authors would like to thank Sandy Pernitzsch/Scigraphix for assistance with Figures 5 and 6.

## Author contributions

Conceptualization, J.V., L.B., J.S.; Investigation, B.M.G., S.G., A.J.W., J.W., S. C., L.C.W., P.S., L.B.; Formal Analysis, B.M.G., W.K.C., S.S., R.S., L.B.; Resources, R.S., J.V., L.B., J.S.; Writing – Original Draft, L.B., J.S.; Writing – Review and Editing, A.J.W., L.B., J.S.; Supervision, L.B., J.S.; Funding Acquisition, J.V., J.S.

