## Supplementary Figures for "Dual RNA-seq provides insight into the biology of the neglected intracellular human pathogen *Orientia tsutsugamushi*"

SUPPLEMENTARY MATERIAL

Bozena Mika-Gospodorz[#a], Suparat Giengkam[#b], Alexander J. Westermann[a,c], Jantana Wongsantichon[b], Willow Kion-Crosby[d], Suthida Chuenklin[b], Loo Chien Wang[e], Piyanate Sunyakumthorn[f], Radoslaw M. Sobota[e], Selvakumar Subbian[h], Jörg Vogel[a,c], Lars Barquist[#*a,g] and Jeanne Salje[#*b,h,i]

Running title: RNA sequencing of *Orientia tsutsugamushi*

Helmholtz Institute for RNA-based Infection Research (HIRI), Helmholtz Centre for Infection Research (HZI), Würzburg, Germany[a]

Mahidol-Oxford Tropical Medicine Research Unit, Faculty of Tropical Medicine, Mahidol University, Bangkok, Thailand[b];

Institute for Molecular Infection Biology (IMIB), University of Würzburg, Würzburg, Germany[c]

Rutgers, the State Univeristy of New Jersey, New Jersey, USA[d]

Functional Proteomics Laboratory, Institute of Molecular and Cell Biology, Agency for Science, Technology and Research (A*STAR), Singapore[e].

Armed Forces Research Institute of Medical Sciences, Bangkok, Thailand[f];

Faculty of Medicine, University of Würzburg, Würzburg, Germany[g];

Public Health Research Institute, Rutgers University, New Jersey, USA[h];

Centre for Tropical Medicine and Global Health, Nuffield Department of Medicine, University of Oxford, Oxford, United Kingdom[i];

[#] these authors contributed equally

**Supp. Fig. 1. Ot_Karp time course.** Confocal microscopy images of Karp strain Ot bacteria in HUVEC cells at 0-7 days post infection. Two representative images are shown at each time point. Blue = DAPI (DNA), Red = Evans blue (host cells), green = Ot labelled with Alexa488-click-methionine.

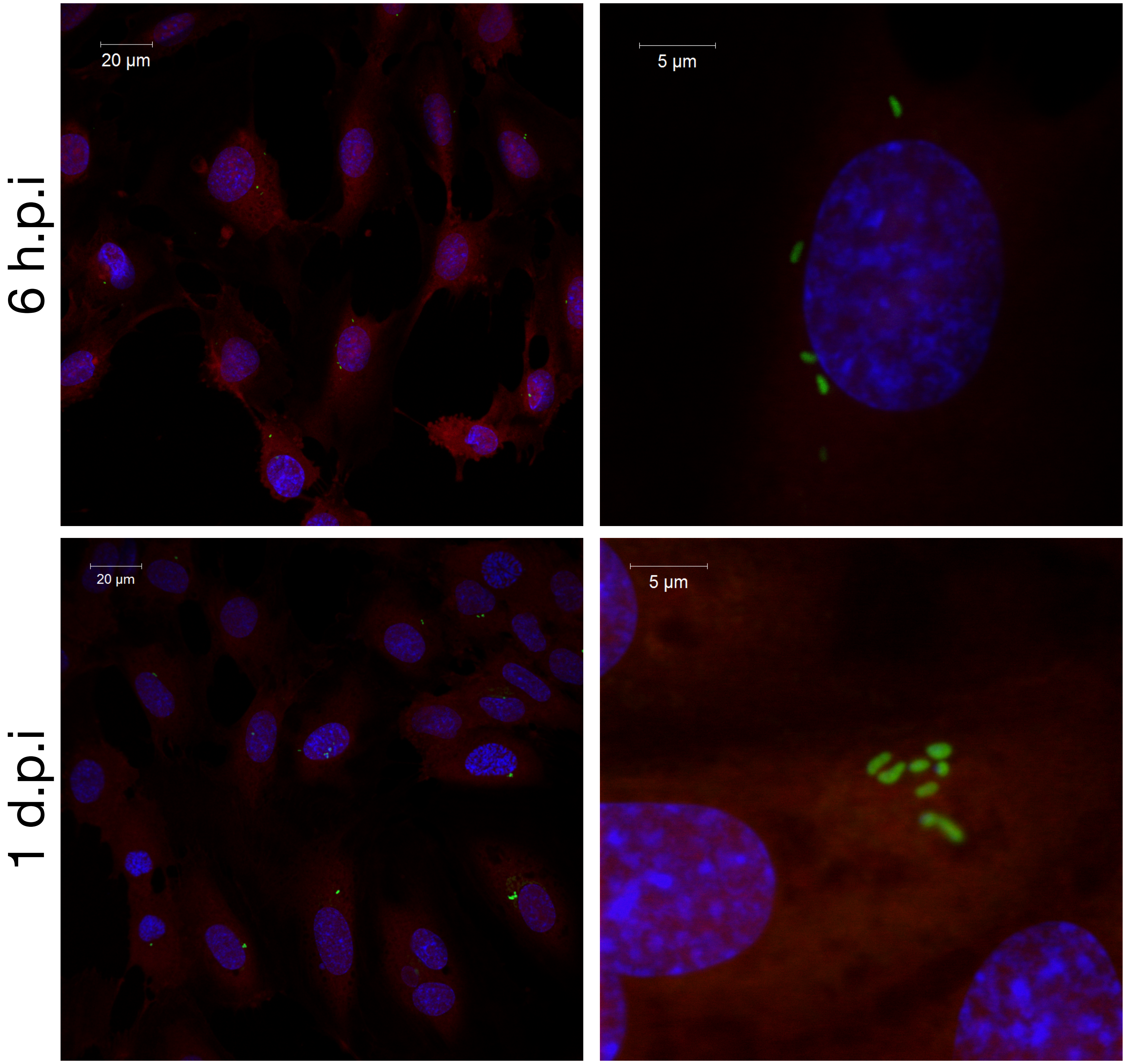

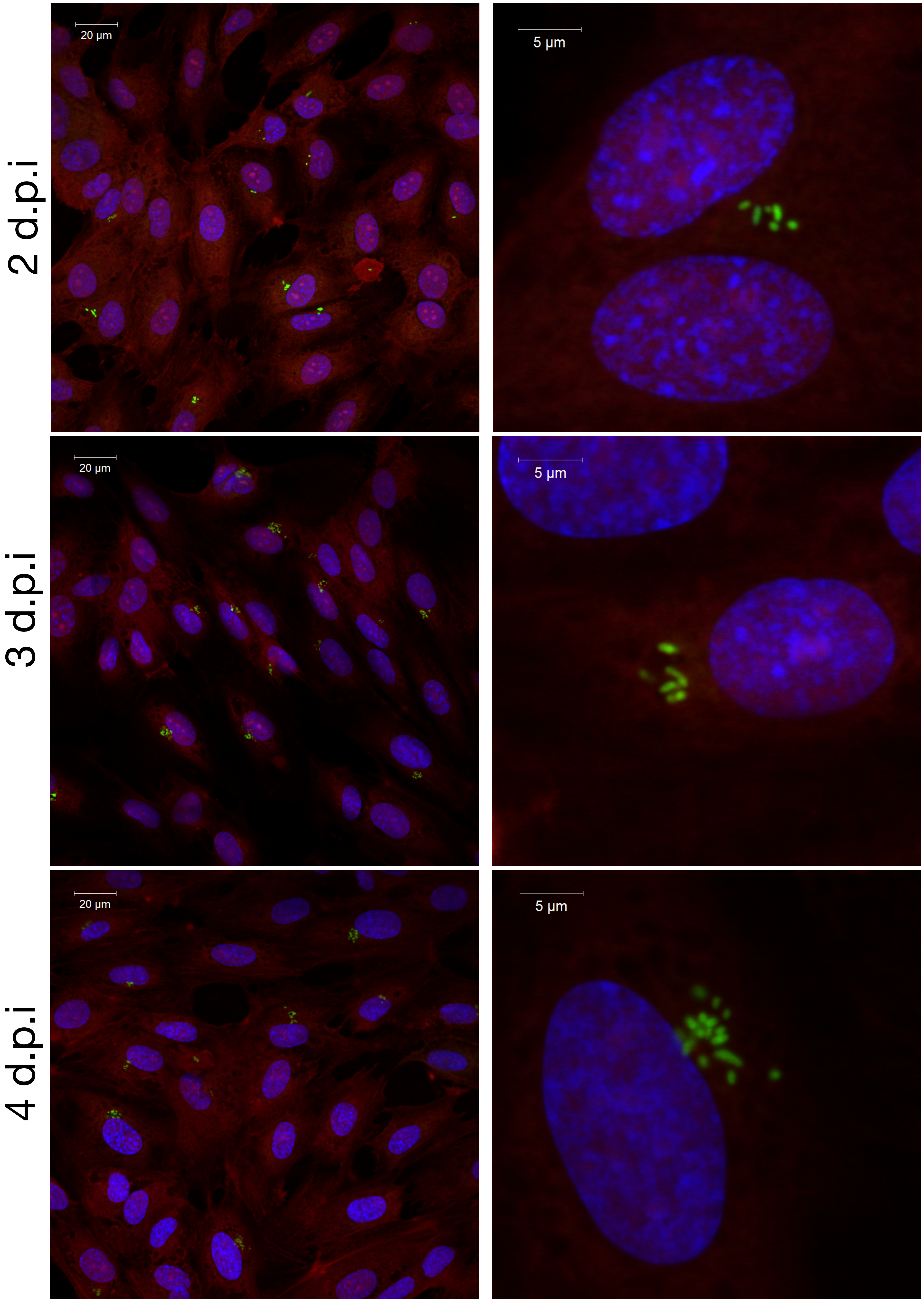
2 d.p.i
3 d.p.i
4 d.p.i
20 µm
5 µm
20 µm
5 µm
20 µm
5 µm

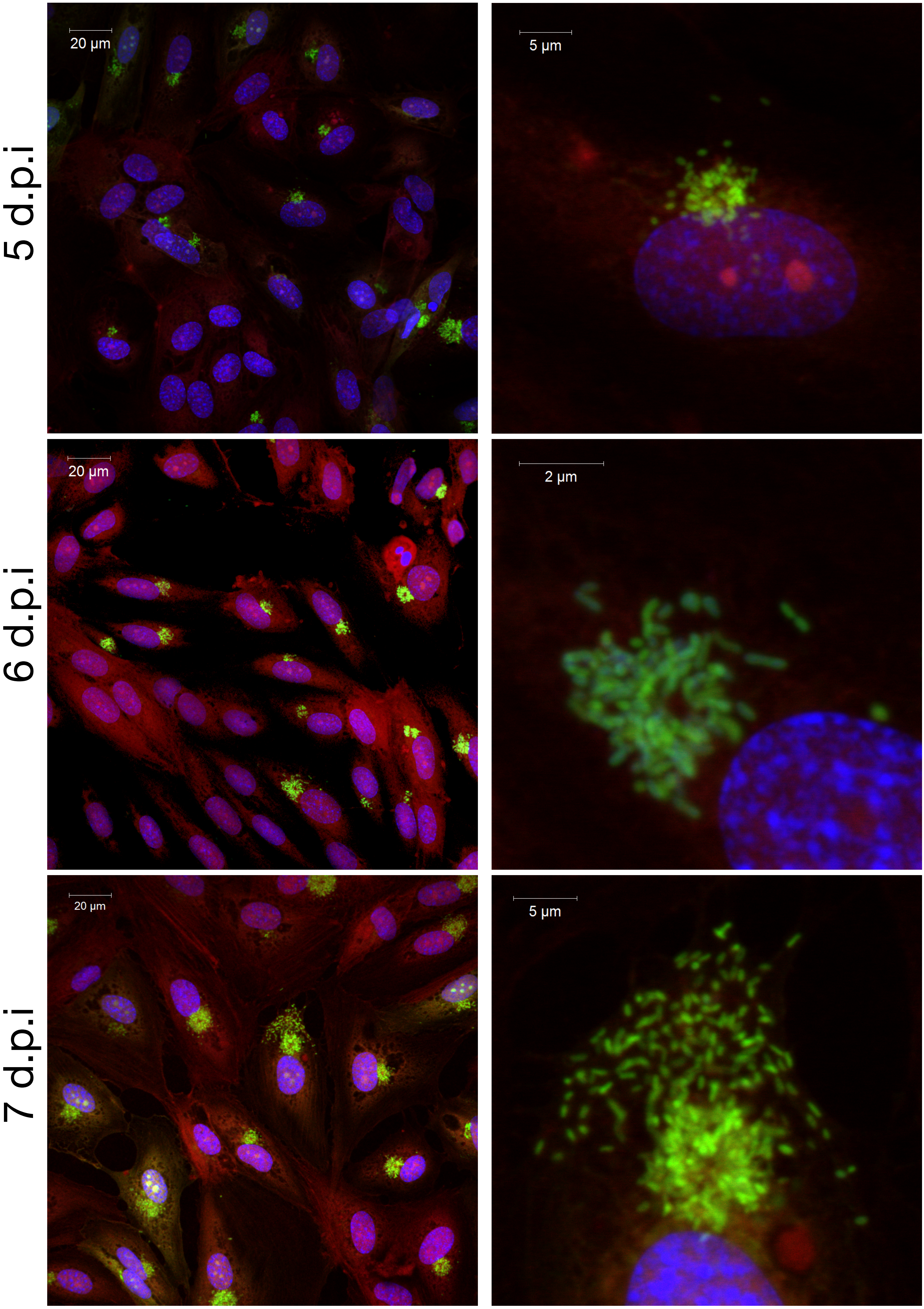
5 d.p.i
20 µm
5 µm
6 d.p.i
20 µm
2 µm
7 d.p.i
20 µm
5 µm

**Supp. Fig. 2. Ot_UT176 time course.** Confocal microscopy images of Ot_UT176 bacteria in HUVEC cells at 0-7 days post infection. Two representative images are shown at each time point. Blue = DAPI (DNA), Red = Evans blue (host cells), green = Ot labelled with Alexa488-click-methionine.

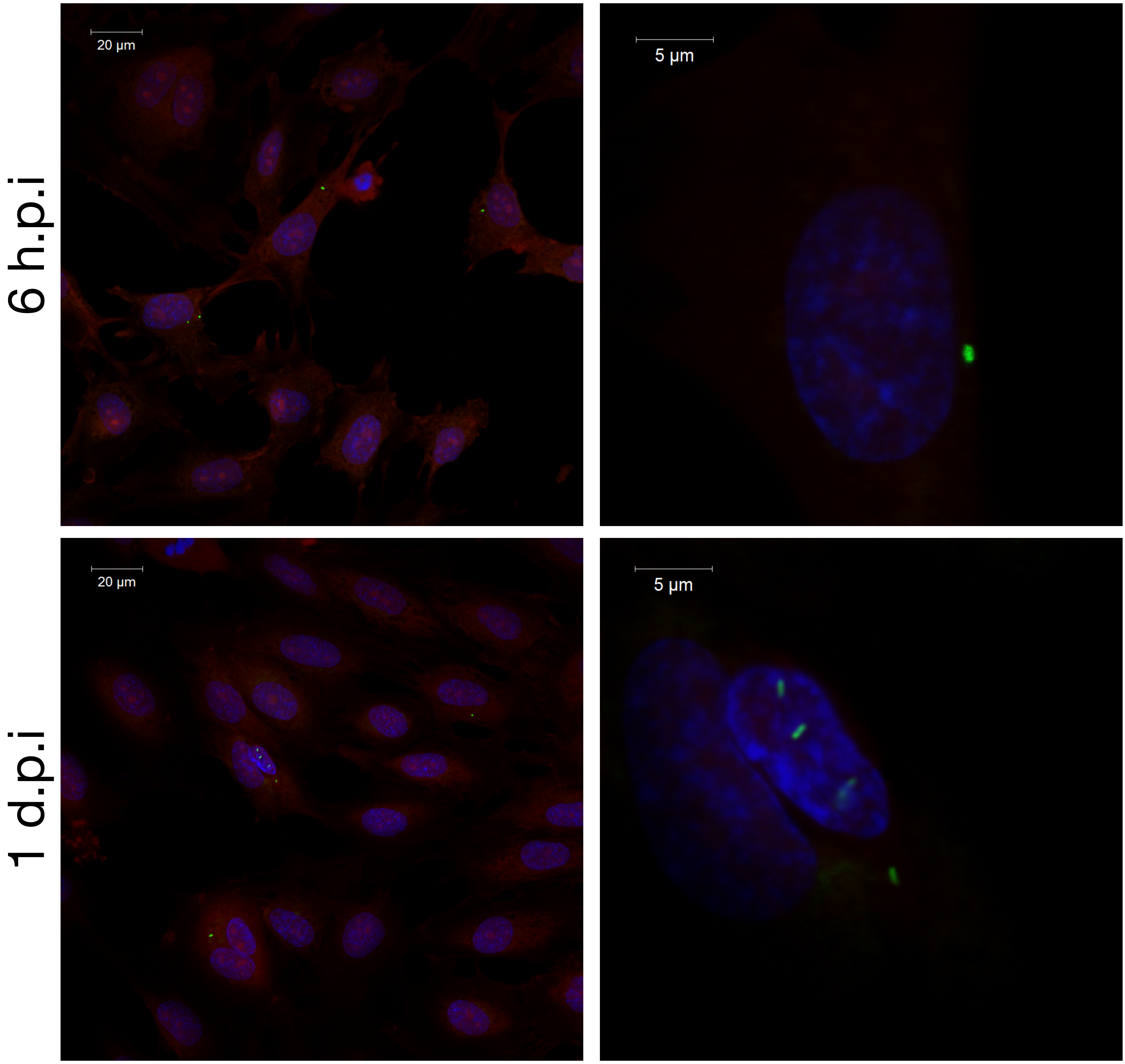

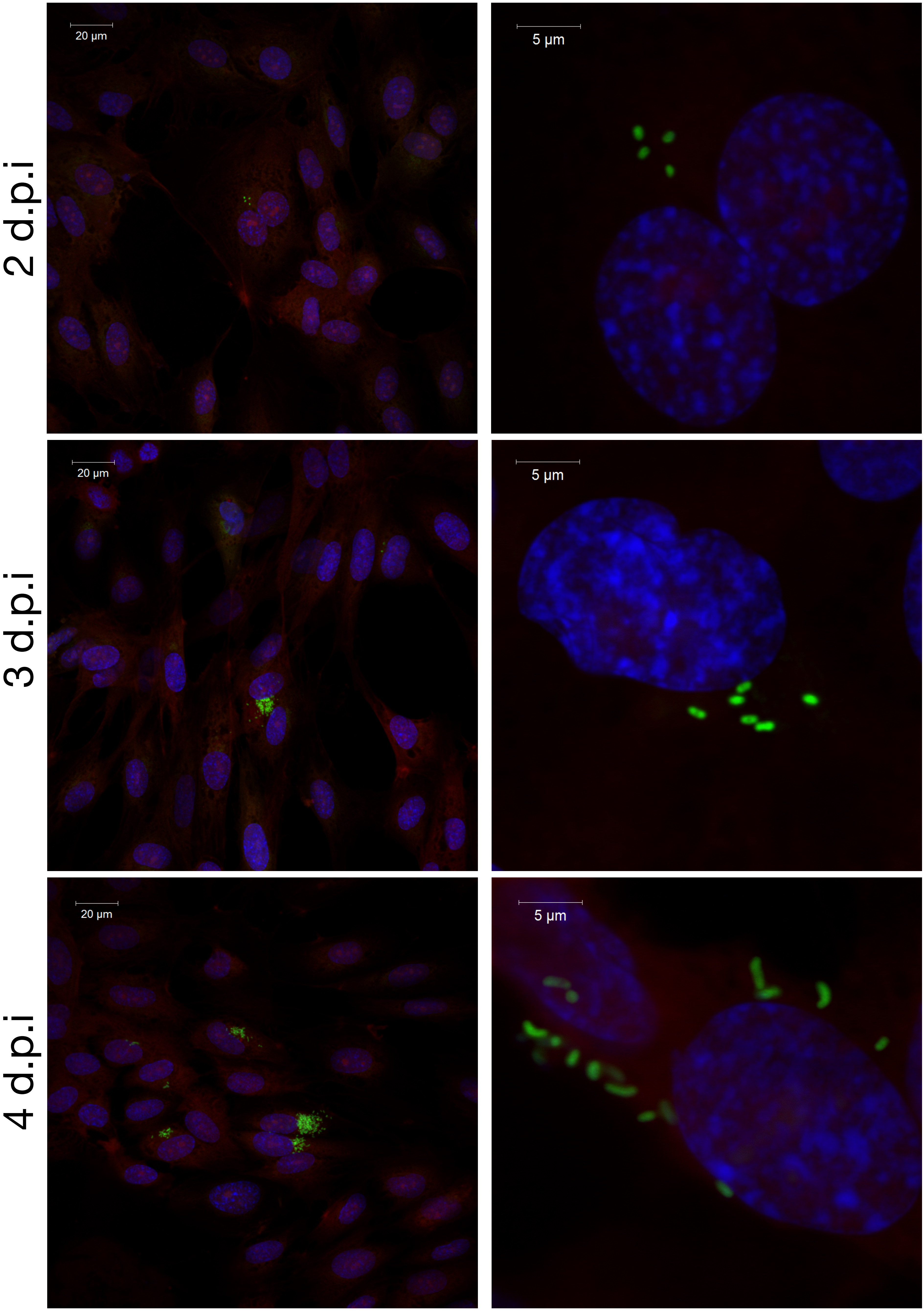
2 d.p.i
3 d.p.i
4 d.p.i
20 μm
5 μm
20 μm
5 μm
20 μm
5 μm

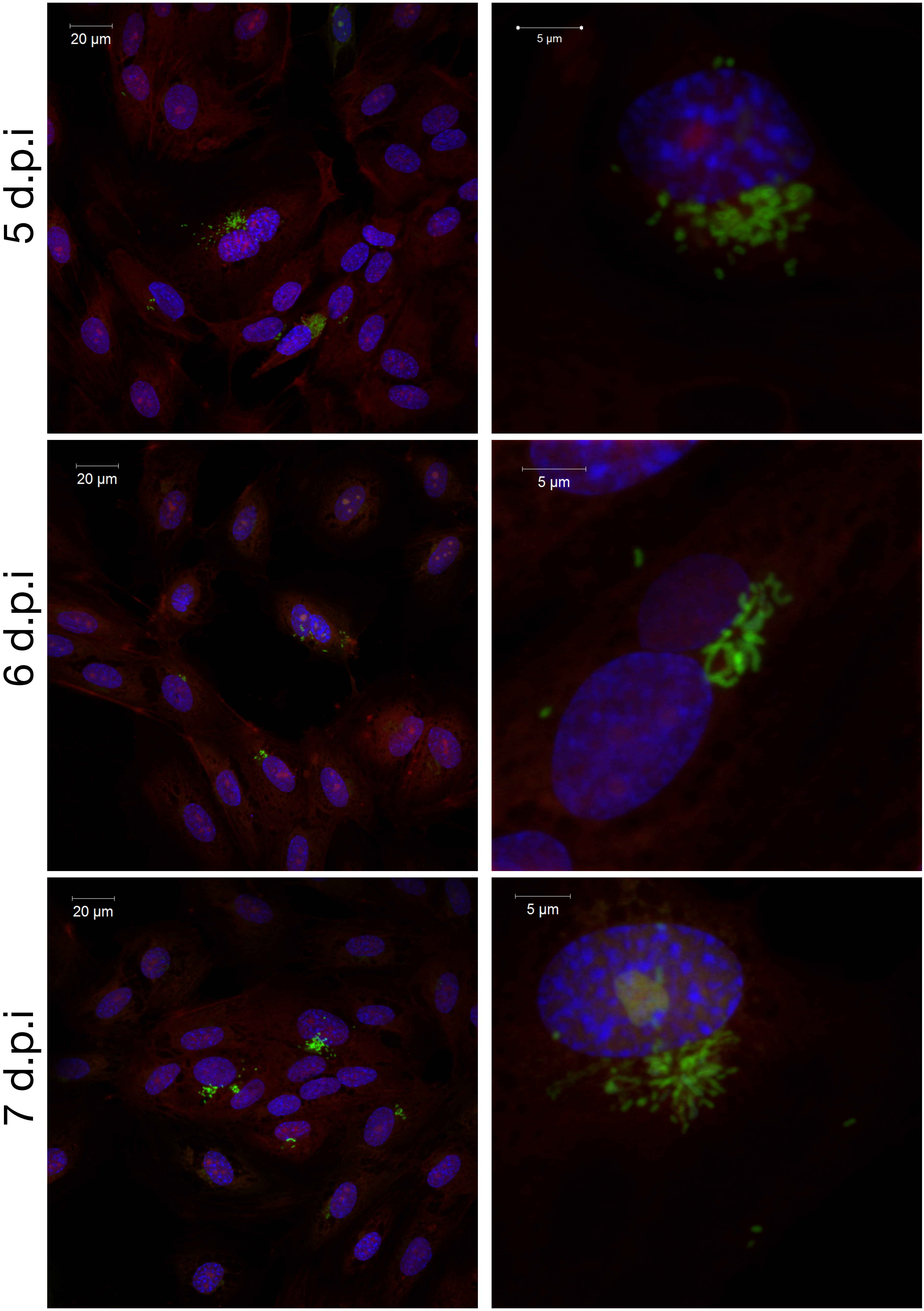
5 d.p.i
20 µm
5 µm
6 d.p.i
20 µm
5 µm
7 d.p.i
20 µm
5 µm

**Supp. Fig. 3. RNA mapping statistics showing ratio of host and Ot RNA in each individual sample.**

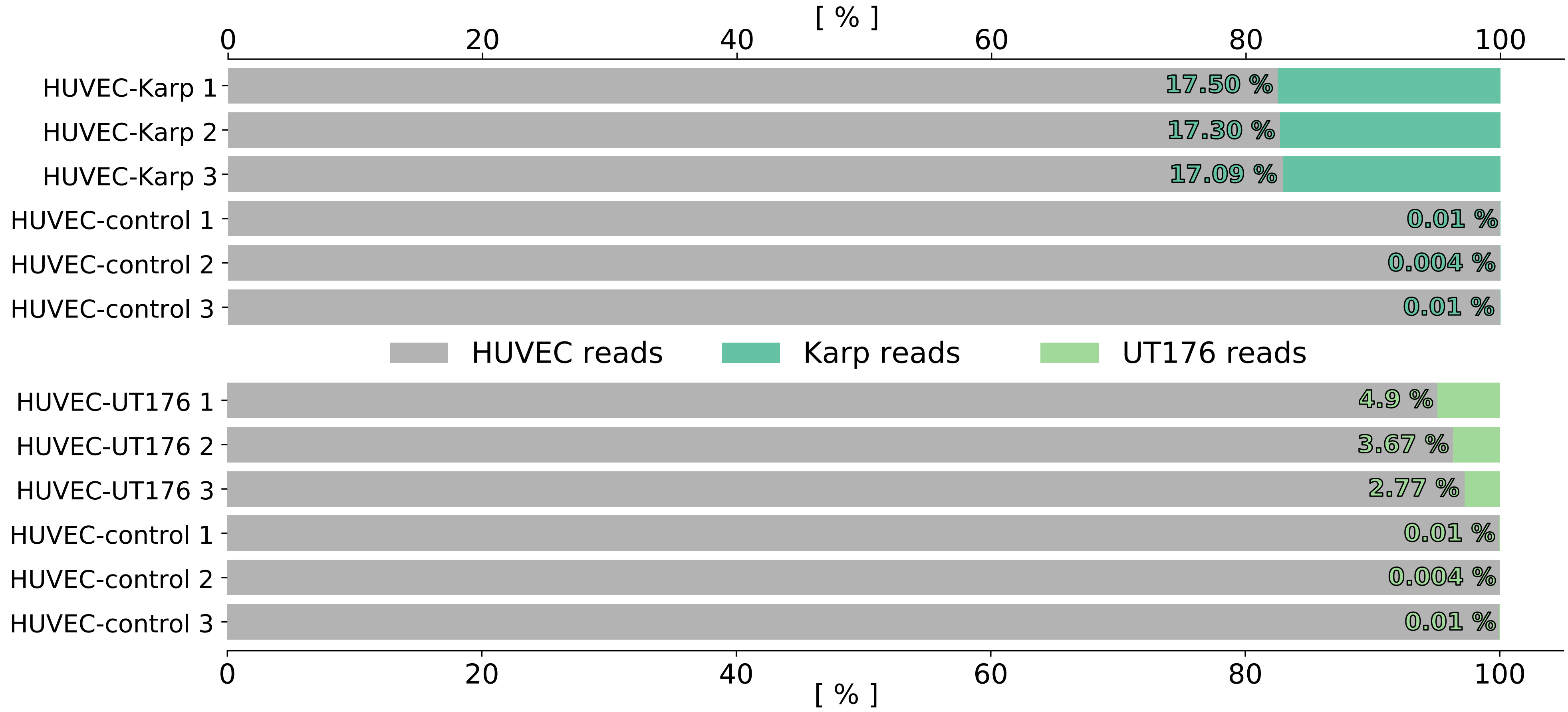

**Supp. Fig. 4. Summary of data reproducibility.** Comparison of replicate log10(TPM) values. Pearson correlation coefficient was calculated for untransformed TPM values.

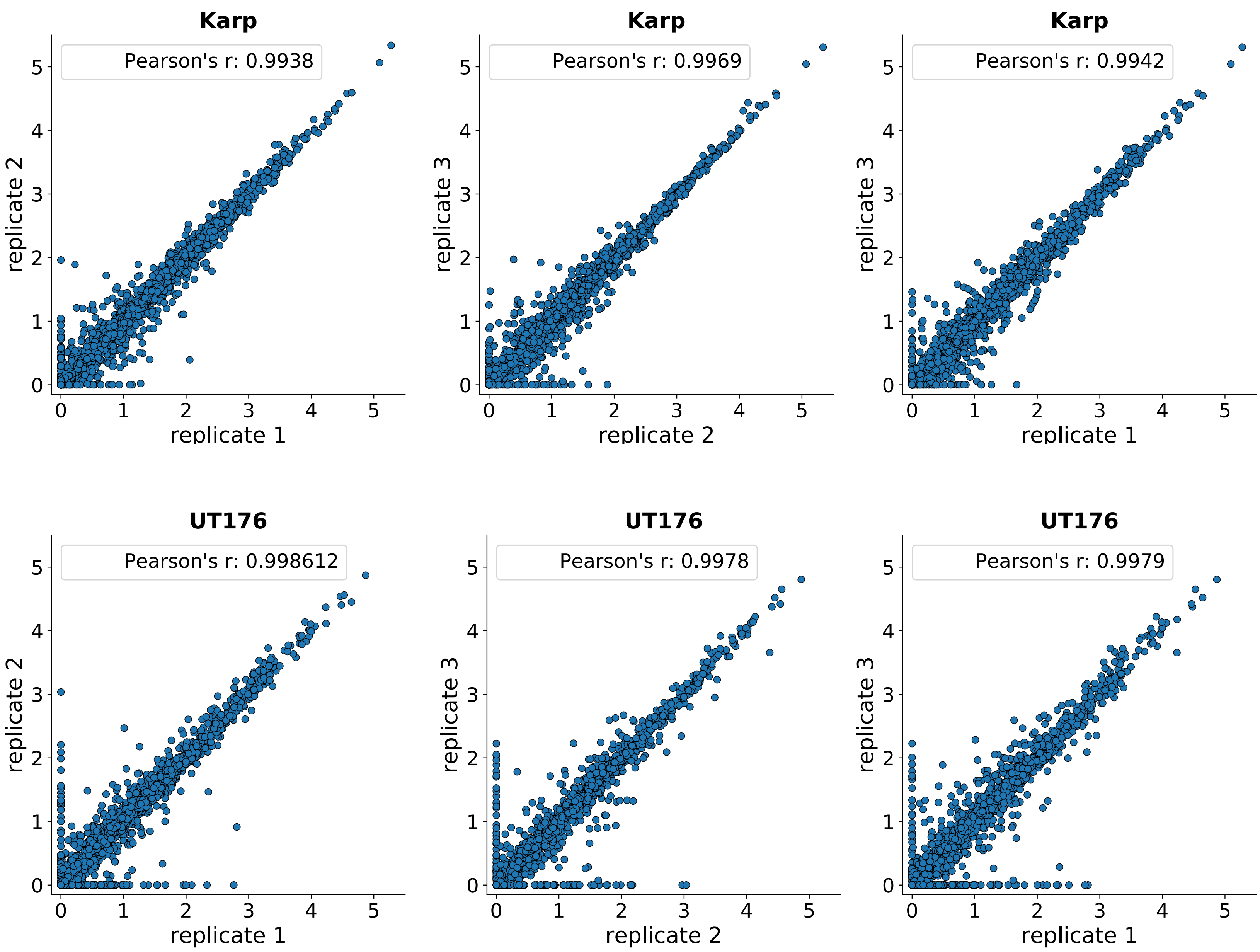

**Supp. Fig. 6. Lengths of conserved operons in Ot**

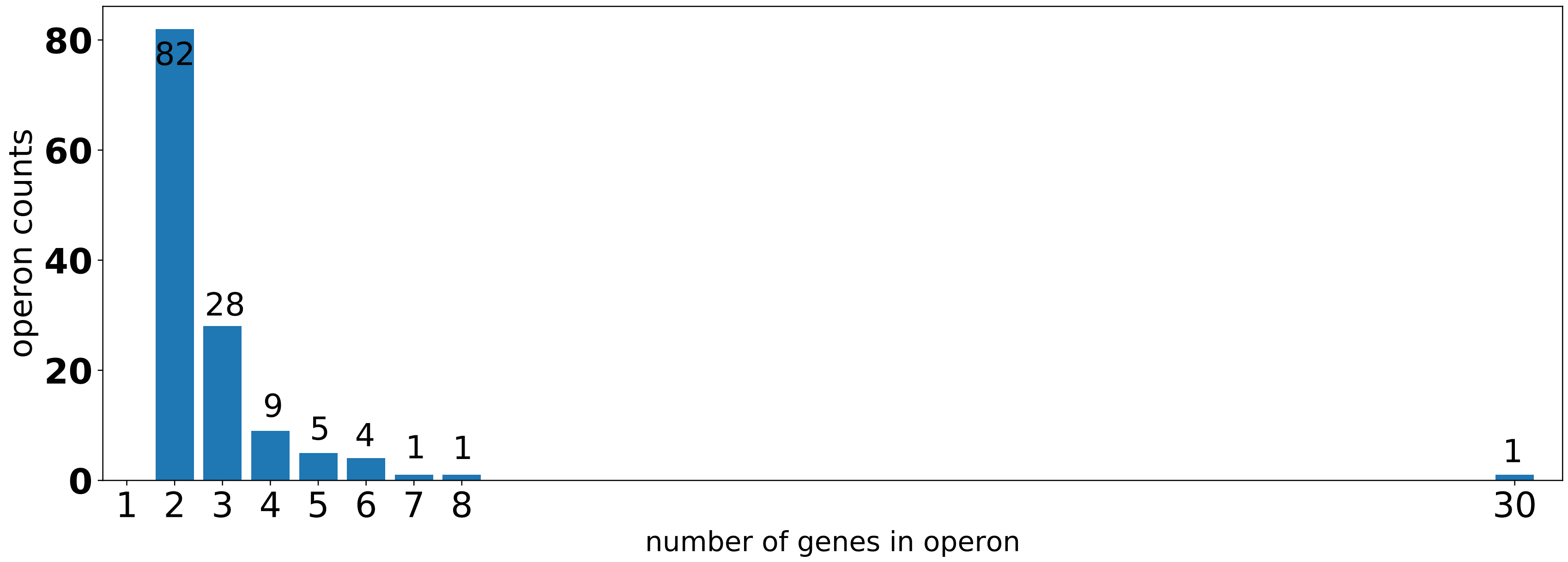

**Supp. Fig. 6. Enrichment of RAGE genes in antisense expression.** Genes were ranked by antisense read counts from highest to lowest, and a hypergeometric p-value was calculated at each position for enrichment of RAGE-annotated genes. Inset plots shows representative annotations for genes with high levels of antisense expression.

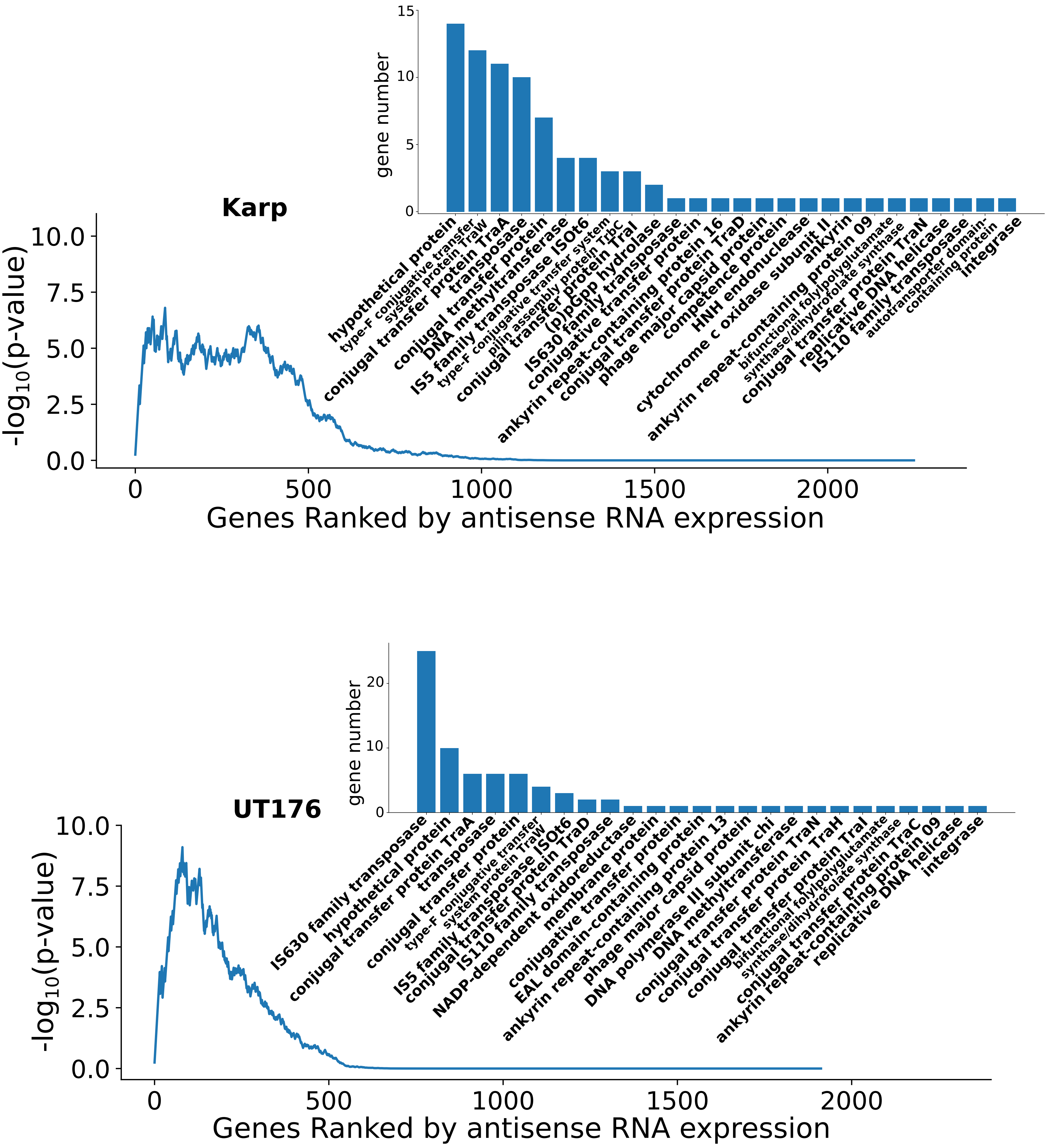

**Supp. Fig. 7. Performance measures of logistic regression models in 500-fold cross validation.** Models incorporating the antisense-sense read ratio (i.e. 2 and 3, see methods) perform better than model 1 based solely on sense expression.

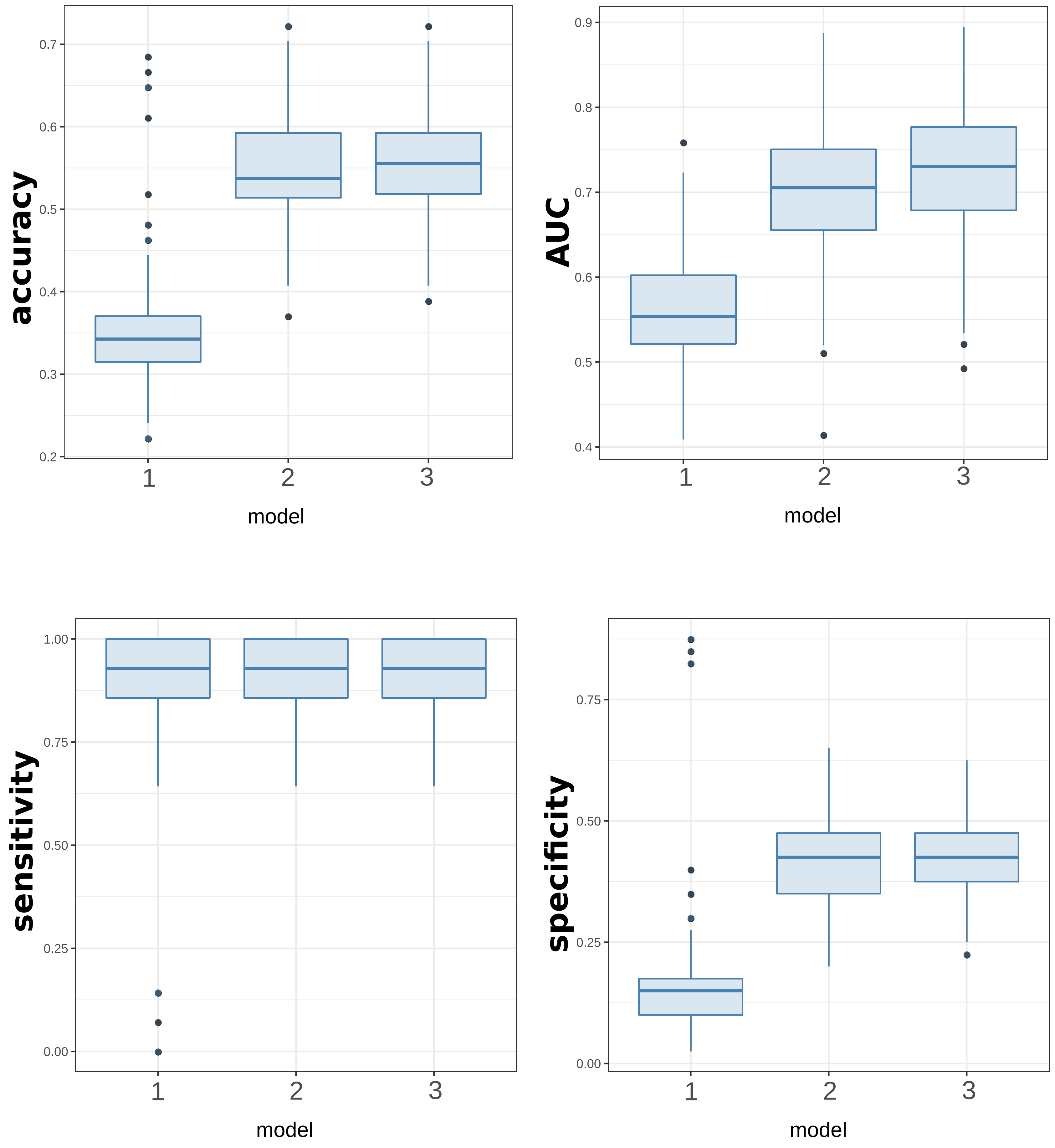

**Supp. Fig. 8. Codon bias in Ot.** A. Genome-wide codon biases in *O. tsutsugamushi, R. typhi, C. crescentus* and *E. coli.* Genome-wide RSCU values are indicated by the green boxes, while the red boxes are the values for 50 randomly chosen individual genes. tRNA gene copy numbers are given in parentheses. B. Relative use of different codons by subsets of genes in Ot (Karp) that are absent in proteomics, detected in proteomics, highly expressed in proteomics, encoded by non-core gene set or encoded by core gene set. Three illustrative examples are shown (Lys, Ile, Ser) with the full set of 20 amino sets given in Supp. Fig. 12. The error bars represent 99.7% confidence intervals and were generated by computing the normalized codon frequencies from 10000 sets of randomly selected genes, with each set being equal in size to the corresponding gene set. C. Overview of relative use of codons to encode each amino acid in the different proteomics gene sets. The bias between groups (yes in green, no in red) indicates whether the different groups use the different possible codons differently. This is defined by a consistent trend between the not detected, detected and highly expressed groups. As an example, Lys in Fig. 4B=no, whilst Ile and Ser=yes. The direction of genome-wide bias indicates whether the genome-wide relative use of the different codon correlates with the presence or absence of cognate tRNA, or not. (positive correlation in green, negative correlation in red, split correlation in yellow). As an example Lys=positive, Ile=negative, Ser=split.

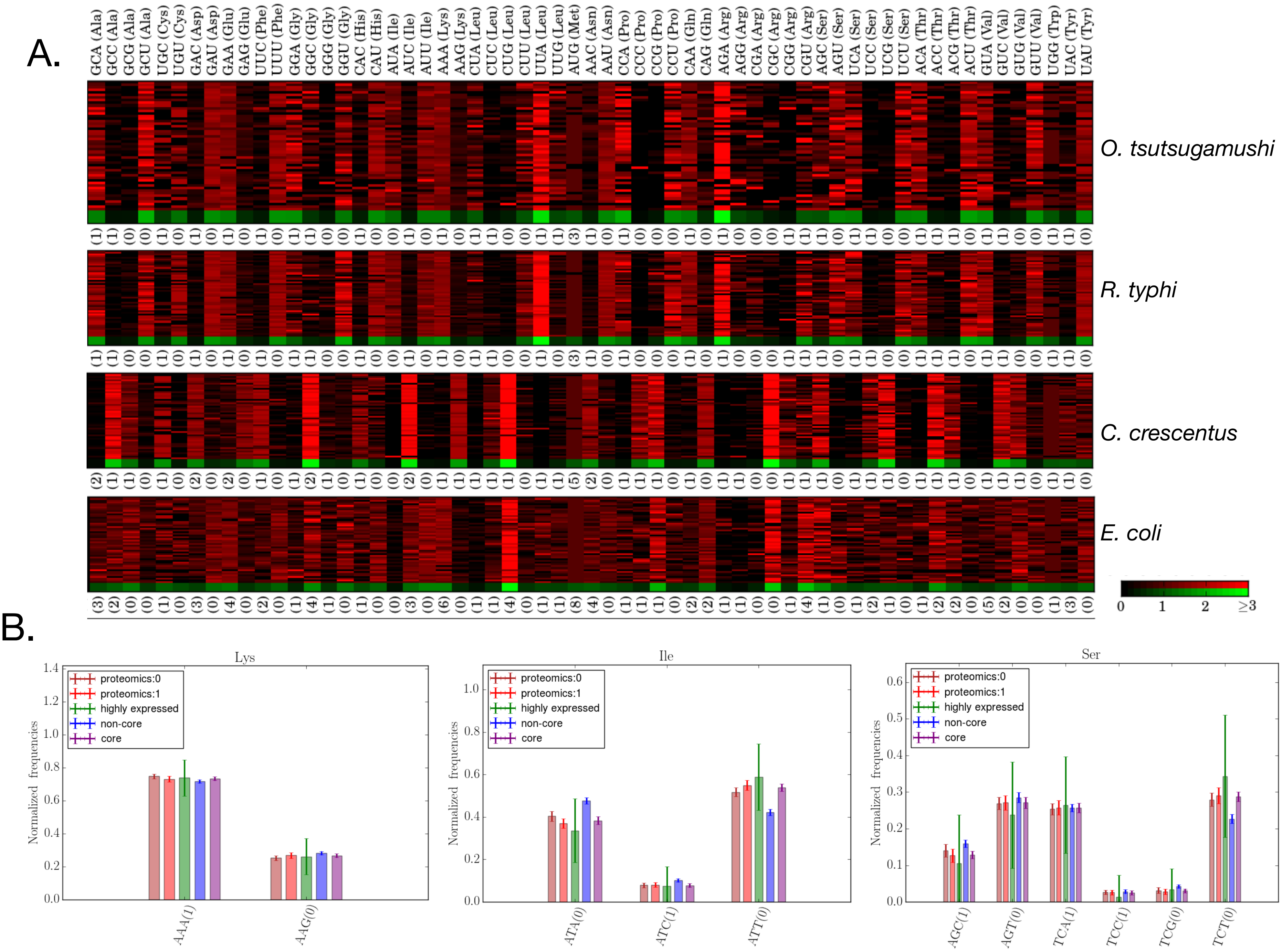

C.

| amino acid | A | C | D | E | F | G | H | I | K | L | M | N | P | Q | R | S | T | V | W | Y |
|---|---|---|---|---|---|---|---|---|---|---|---|---|---|---|---|---|---|---|---|---|
| number of codons | 4 | 2 | 2 | 2 | 2 | 4 | 2 | 3 | 2 | 6 | 1 | 2 | 4 | 2 | 6 | 6 | 4 | 4 | 1 | 2 |
| bias between groups | Y | Y | N | N | N | Y | N | Y | N | N | - | N | Y | N | Y | Y | Y | Y | - | N |
| direction of genome-wide bias | S | N | N | P | N | S | N | N | P | S | - | N | S | P | S | S | S | S | - | N |

**Supp. Fig. 9. Codon biases between different groups of OT_Karp genes**

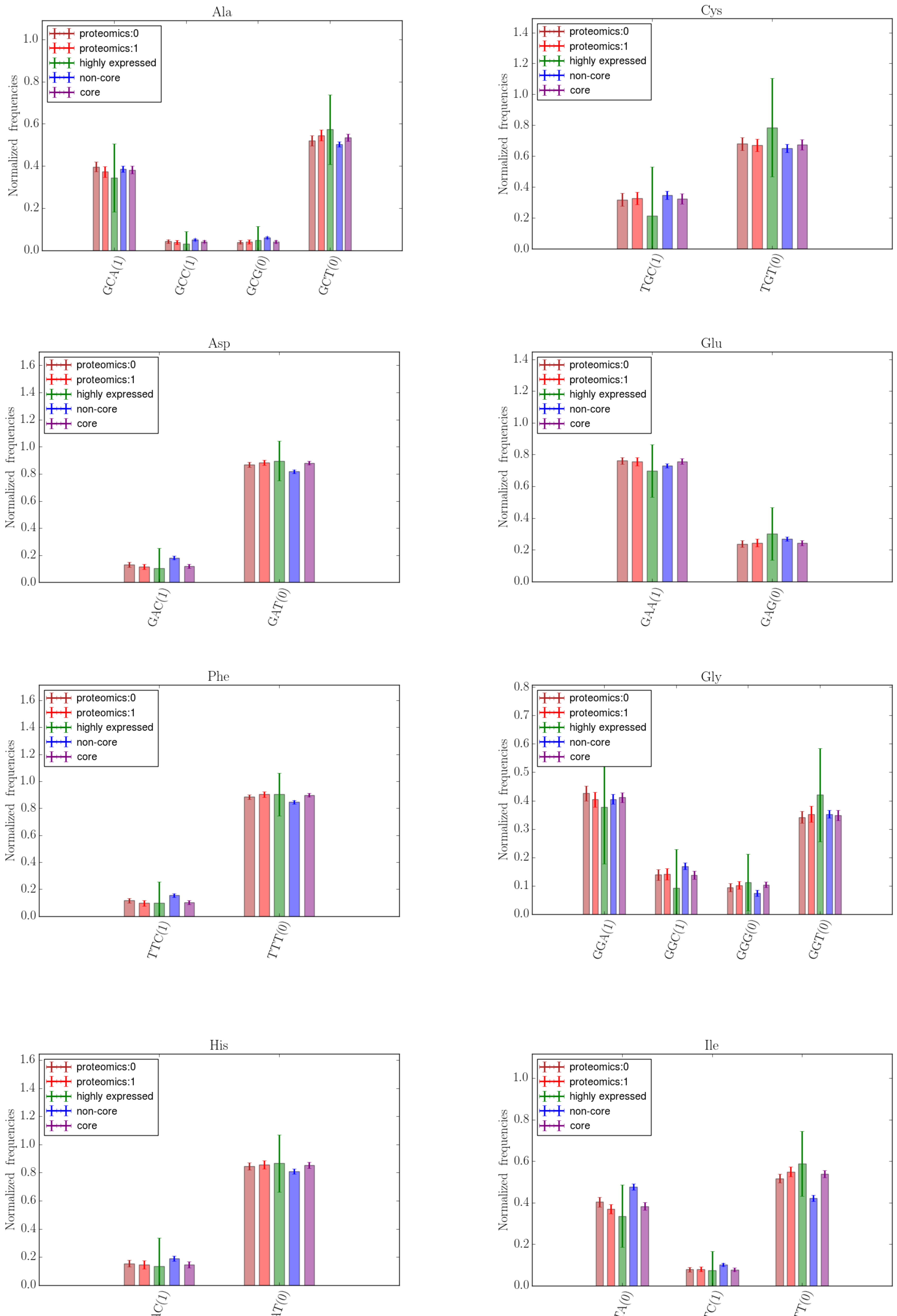

Supp. Fig. 9. cont.

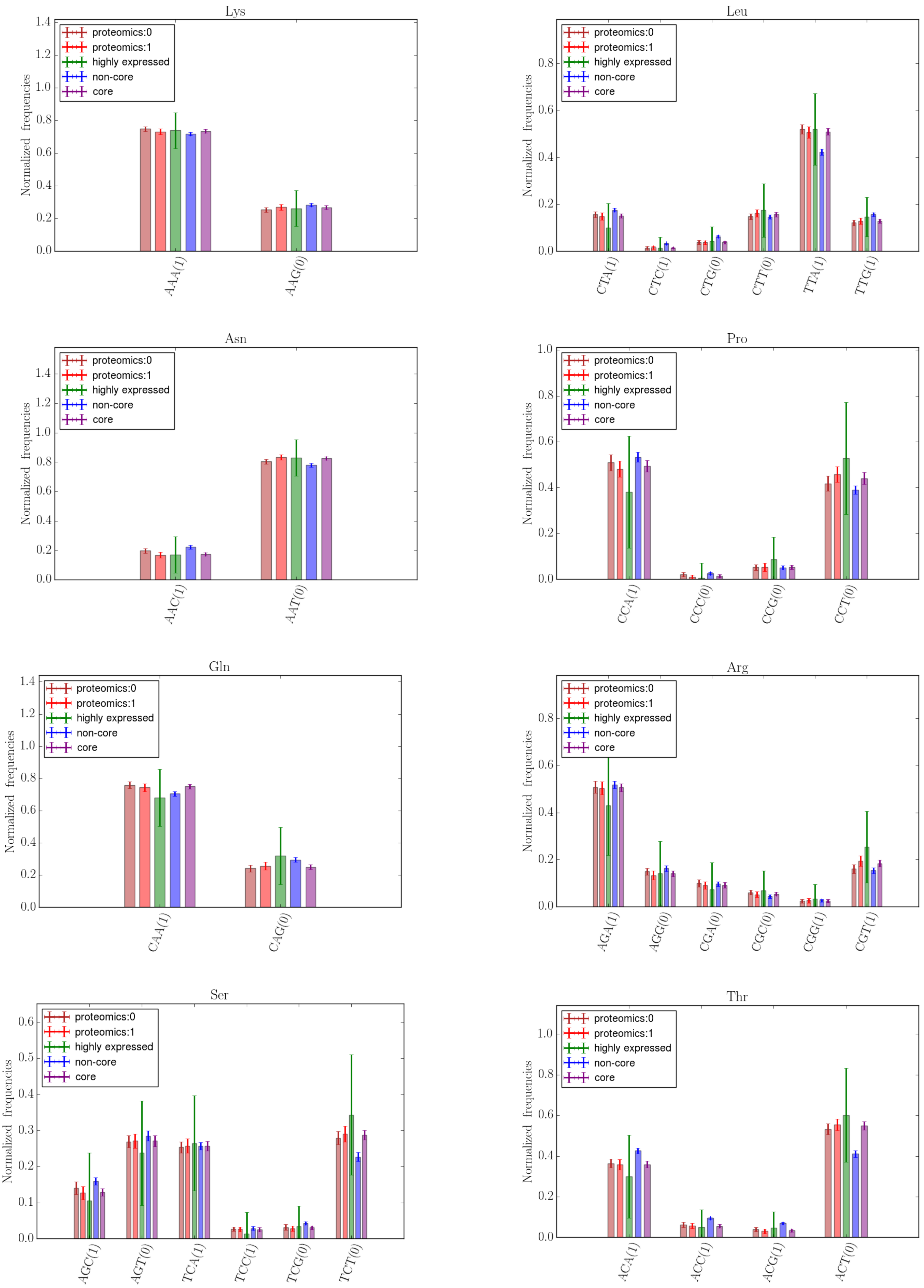

Supp. Fig. 9. cont.

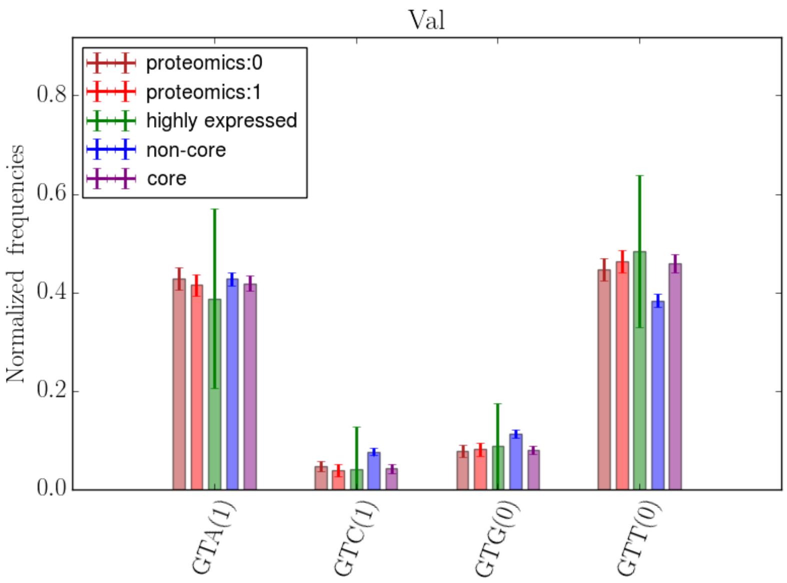

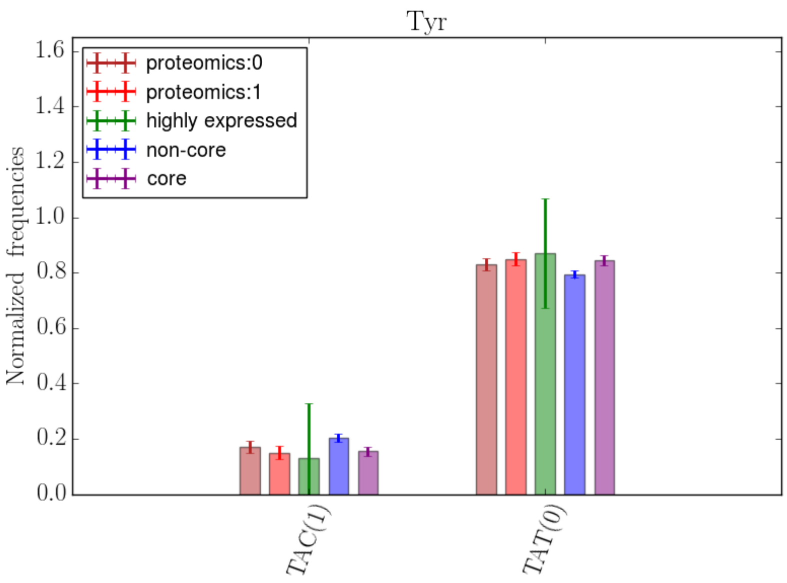

**Supp. Fig. 10. Volcano plots showing differential expression of host genes in HUVEC cells infected with Ot_Karp and Ot_UT176**

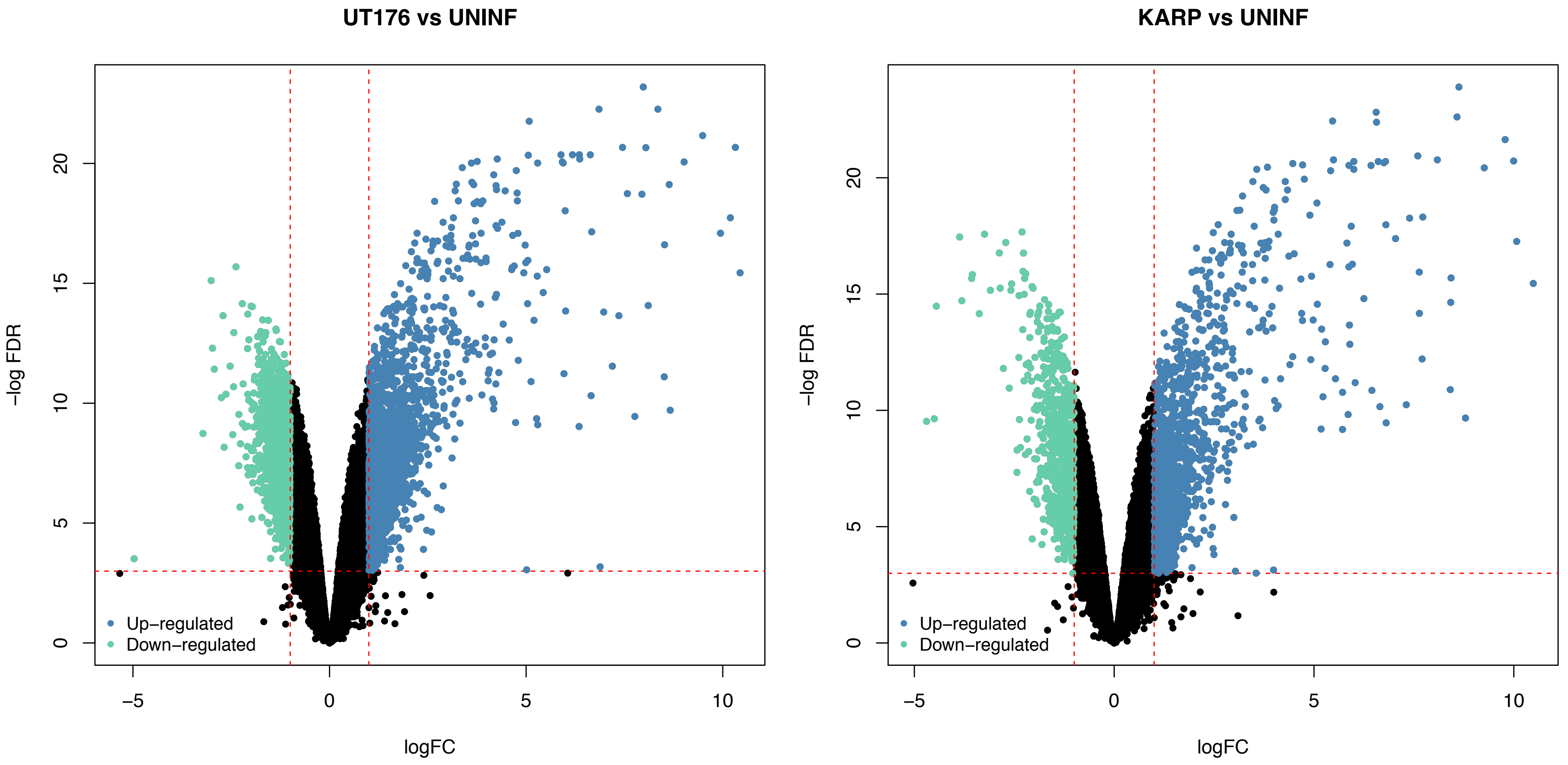

**Supp. Fig. 11. Differential cytokine and chemokine gene expression pattern in Ot_KARP or Ot_UT176 infected HUVEC cells measured by RNAseq**

| Symbol | KARP | UT178 |
|---|---|---|
| *CXCL10* | 8.4 | 10.2 |
| *IFNB1* | 8.8 | 8.7 |
| *CXCL6* | 0.0 | 5.3 |
| *CCL5* | 4.1 | 4.8 |
| *CXCL11* | 4.4 | 4.7 |
| *CXCL5* | 0.0 | 4.2 |
| *CX3CL1* | 0.9 | 3.6 |
| *CCL20* | -1.2 | 3.6 |
| *IL7R* | 3.8 | 3.5 |
| *TNFSF10* | 3.6 | 3.4 |
| *CXCL1* | 0.0 | 3.2 |
| *IL1RL1* | 0.5 | 3.0 |
| *CXCL8* | -2.0 | 2.9 |
| *CXCL3* | 0.0 | 2.9 |
| *IL18R1* | 0.5 | 2.7 |
| *TNFAIP3* | 1.2 | 2.7 |
| *IL34* | 0.0 | 2.6 |
| *IL6* | 0.0 | 2.5 |
| *IL3RA* | 2.1 | 2.2 |
| *IL15RA* | 2.1 | 2.1 |
| *IL1A* | 0.0 | 2.1 |
| *TNFSF13B* | 1.7 | 2.1 |
| *IL13RA2* | 5.9 | 2.0 |
| *CXCL2* | -0.5 | 1.9 |
| *IL15* | 1.1 | 1.7 |
| *TNFRSF9* | 0.0 | 1.7 |
| *IL1R1* | 0.0 | 1.7 |
| *TNFAIP2* | 0.6 | 1.6 |
| *IL33* | 6.6 | 1.5 |
| *TNFSF18* | 0.0 | 1.4 |
| *IL12A* | 0.0 | 1.3 |
| *TNFRSF1B* | 0.9 | 1.3 |
| *IL18BP* | 1.3 | 1.3 |
| *CCL2* | -1.5 | 1.1 |
| *TNFRSF25* | 1.2 | 1.0 |
| *IL11RA* | 0.7 | 0.9 |

| Symbol | KARP | UT178 |
|---|---|---|
| *TNFAIP6* | 0.0 | 0.9 |
| *TNFAIP8* | 0.0 | 0.9 |
| *IFNAR2* | 0.8 | 0.8 |
| *IL20RB* | 0.0 | 0.8 |
| *CXCL16* | 1.1 | 0.8 |
| *TNFRSF11A* | 1.4 | 0.8 |
| *IL6R* | 1.1 | 0.8 |
| *TNFRSF14* | 0.0 | 0.7 |
| *IL4R* | 0.0 | 0.6 |
| *IL4I1* | 0.0 | 0.6 |
| *IFNGR2* | -0.3 | 0.6 |
| *TNFAIP8L1* | 0.0 | 0.5 |
| *CXCR4* | 0.0 | 0.5 |
| *IL1RAP* | 0.0 | 0.5 |
| *IL1RAPL2* | 1.8 | 0.0 |
| *TNFAIP8L3* | 1.3 | 0.0 |
| *IL1RAPL1* | 1.0 | 0.0 |
| *TNFRSF10A* | 0.8 | 0.0 |
| *TNFRSF6B* | 0.7 | 0.0 |
| *IL6ST* | -0.4 | 0.0 |
| *TNFRSF12A* | -0.4 | 0.0 |
| *IL17RA* | -0.4 | 0.0 |
| *TNFRSF10B* | -0.5 | 0.0 |
| *IL6STP1* | -0.6 | 0.0 |
| *IL17D* | -0.6 | 0.0 |
| *TNFRSF19* | -0.8 | 0.0 |
| *TNFSF15* | -2.6 | 0.0 |
| *TNFRSF1A* | 0.4 | -0.3 |
| *TNFRSF10C* | -0.4 | -0.4 |
| *TNFSF12* | 0.0 | -0.5 |
| *IL13RA1* | -1.0 | -0.5 |
| *IL32* | -1.1 | -0.7 |
| *TNFRSF21* | 0.3 | -0.7 |
| *TNFSF4* | 0.9 | -1.4 |
| *TNFRSF10D* | -1.6 | -1.7 |

**Supp Fig 12. Differential analysis of host genes. qRT-PCR of host genes in HUVEC cells infected with Ot_Karp or Ot_UT176**

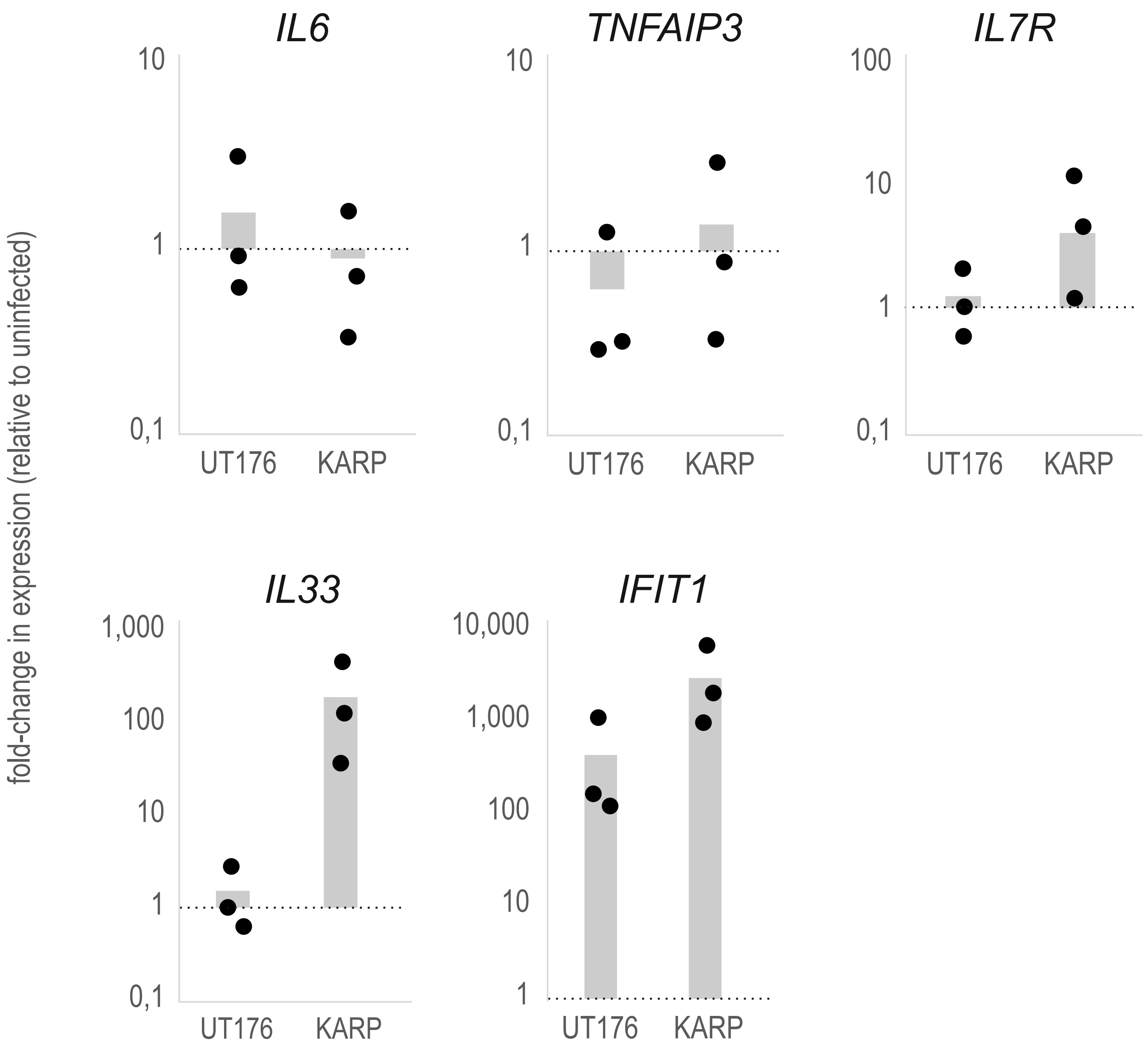

**Supp Fig 13. Summary of primers and probes used in this study**

| Northern blot probes | name | sequence (5'->3') | length [nt] | description | reference |
|---|---|---|---|---|---|
| | AWO-009 | TACCTCTATTCTTAA TAAAACTTATTGCC | 29 | antisense Northern probe against Orientia tmRNA (5') | this study |
| | AWO-010 | TGATTTCCTTAAGCT GCTAATG | 22 | antisense Northern probe against Orientia tmRNA (3') | this study |
| | AWO-011 | GGACTTTCCTCACA AATCTAT | 21 | antisense Northern probe against Orientia RNaseP RNA | this study |
| | AWO-013 | GTTGATGCCTACGC CAGTTA | 20 | antisense Northern probe against Orientia SRP RNA | this study |
| | AWO-022 | CTCTCCCATGTTTA AACATA | 20 | antisense Northern probe against Orientia 5S rRNA | this study |
| | JVO-7672 | ATATGGAACGCTTC ACGAATTTG | 23 | antisense Northern probe against human U6 snRNA | PMID:26789254 |
| **qRT-PCR primers** | **name** | **sequence (5'->3')** | **length [nt]** | **description** | **reference** |
| | JVO-8896 | TCGGTACATCCTCG ACGG | 18 | sense qPCR primer against human IL6 mRNA | this study |
| | JVO-8897 | TGTTTTCTGCCAGT GCCTC | 19 | antisense qPCR primer against human IL6 mRNA | this study |
| | JVO-14331 | GCTGTGAAGATACG GGAGAGAAC | 23 | sense qPCR primer against human TNFAIP3 mRNA | this study |
| | JVO-14332 | CCTGGATGTTTCTG TCGATGAG | 22 | antisense qPCR primer against human TNFAIP3 mRNA | this study |
| | JVO-9476 | AAGTGGCTATGCTC AAAATG | 20 | sense qPCR against human IL7R mRNA | PMID:21307942 |
| | JVO-9477 | TTCAGGCACTTTAC CTCCAC | 20 | antisense qPCR against human IL7R mRNA | PMID:21307942 |
| | AWO-007 | CAAACGATAGGCTC AAACACT | 22 | sense qRT-PCR oligo for human IL33 mRNA | this study |
| | AWO-008 | TGAGTGTTGCCTAA GACATC | 20 | antisense qRT-PCR oligo for human IL33 mRNA | this study |
| | JVO-13531 | ATGCAGGAAGAACA TGACAACC | 22 | sense qRT-PCR oligo for human IFIT1 mRNA | this study |
| | JVO-13532 | TCTGGACACTCCAT TCTATAGCG | 23 | antisense qRT-PCR oligo for human IFIT1 mRNA | this study |
| | JVO-7673 | GCTTCGGCAGCAC ATATACTAAAAT | 25 | sense qPCR primer against human U6 snRNA | PMID:26789254 |
| | JVO-7672 | ATATGGAACGCTTC ACGAATTTG | 23 | antisense qPCR primer against human U6 snRNA | PMID:26789254 |

**Supp Fig 14. Karp and UT176 lead to up-regulation of distinct networks in HUVEC cells**

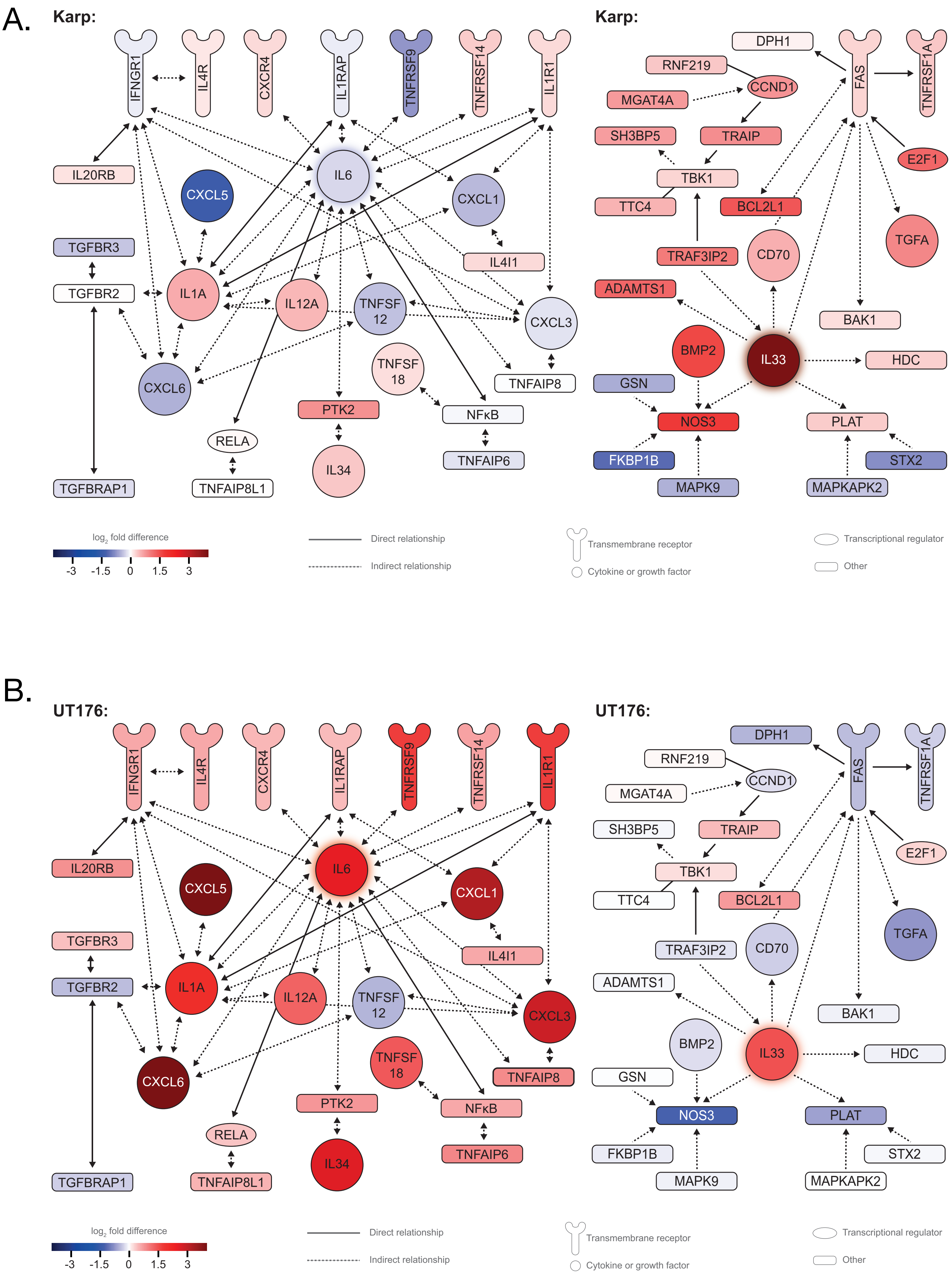

**Supp Fig 15. Differential regulation of inflammatory pathways by UT176 and Karp.** A. shows expression of NFkB pathway genes in UT176- and Karp-infected host cells. B. shows expression of host genes associated with NOS2 production. Red indicates increased expression relative to uninfected cells, blue indicates decreased expression.

A.

| Symbol | Entrez Gene Name | Expr Log Ratio (UT176 vs | Expr Log Ratio (KARP vs | Location | Gene ID-Huma | Gene ID-Mous |
|---|---|---|---|---|---|---|
| *Cot* | mitogen-activated protein kinase kinase kinase 8 | 1.70782 | 0.144809 | Cytoplasm | 1326 | 26410 |
| *RelB* | RELB proto-oncogene, NF-kB subunit | 1.28783 | -0.03004 | Nucleus | 5971 | 19698 |
| *MEKK1* | mitogen-activated protein kinase kinase kinase 1 | 0.92341 | -0.0359 | Cytoplasm | 4214 | 26401 |
| *JNK1* | mitogen-activated protein kinase 8 | 0.817456 | 0.423134 | Cytoplasm | 5599 | 26419 |
| *NF-ϰB2 p100* | nuclear factor kappa B subunit 2 | 0.766144 | -0.39373 | Nucleus | 4791 | 18034 |
| *NF-ϰB1* | nuclear factor kappa B subunit 1 | 0.599586 | -0.03203 | Nucleus | 4790 | 18033 |
| *β-TrCP* | beta-transducin repeat containing E3 ubiquitin protein ligase | 0.465374 | 0.146903 | Cytoplasm | 8945 | 12234 |
| *MALT1* | MALT1 paracaspase | 0.404342 | 0.115903 | Cytoplasm | 10892 | 240354 |
| *p65/RelA* | RELA proto-oncogene, NF-kB subunit | 0.400752 | 0.054767 | Nucleus | 5970 | 19697 |
| *LTBR* | lymphotoxin beta receptor | -0.3528 | -0.22677 | Plasma Membrane | 4055 | 17000 |
| *IKKα* | conserved helix-loop-helix ubiquitous kinase | -0.39103 | 0.0463 | Cytoplasm | 1147 | 12675 |
| *Bcl10* | B cell CLL/lymphoma 10 | -0.67364 | -0.04115 | Cytoplasm | 8915 | 12042 |

Expression of NFkB pathway genes

B.

| Sym bol | Entrez Gene Name | Expr Log Ratio (UT176 vs UnInf) | Expr Log Ratio (KARP vs UnInf) | Locatio n | Gene ID-Human | Gene ID-Mouse |
|---|---|---|---|---|---|---|
| *PPA Rα* | peroxisome proliferator activated receptor alpha | 0.955636 | 0.256633 | Nucleus | 5465 | 19013 |
| *SIR Pα* | signal regulatory protein alpha | 0.855516 | 0.231361 | Plasma Membra ne | 140885 | 19261 |
| *ME K1* | mitogen-activated protein kinase kinase 1 | 0.726498 | 0.240664 | Cytoplas m | 5604 | 26395 |
| *CBP* | CREB binding protein | 0.365126 | 0.187724 | Nucleus | 1387 | 12914 |

Expression of genes involved in NOS2 production

**Supp Fig 16. Differential activation of inflammatory pathways by UT176 and Karp.** A. shows expression of host genes associated with differentiation of mononuclear leukocytes B. shows expression of host genes associated with leukocyte proliferation. The predicted effect on leukocyte differentiation/proliferation is shown, based on the gene expression in response to UT176 infection. Red indicates increased expression relative to uninfected cells, blue indicates decreased expression.

### A. Expression of host genes associated with differentiation of mononuclear leukocytes

| ID | Genes in dataset | Expr Log Ratio (UT176 vs Uninf) | Expr Log Ratio (KARP vs Uninf) | Prediction in UT176 (based on measurement direction) |
|---|---|---|---|---|
| ENSG00000171855.6 | *IFNB1* | 8.664 | 8.790328 | Increased |
| ENSG00000164400.5 | *CSF2* | 7.764 | -1.667 | Increased |
| ENSG00000184979.9 | *USP18* | 5.078 | 5.464608 | Increased |
| ENSG00000108342.12 | *CSF3* | 5.037 | 2.804455 | Increased |
| ENSG00000114251.13 | *WNT5A* | 4.976 | 3.61382 | Increased |
| ENSG00000271503.5 | *CCL5* | 4.801 | 4.102429 | Increased |
| ENSG00000137462.6 | *TLR2* | 4.416 | -0.21786 | Increased |
| ENSG00000163735.6 | *CXCL5* | 4.174 | -1.41573 | Increased |
| ENSG00000057657.14 | *PRDM1* | 3.961 | 1.080978 | Increased |
| ENSG00000128917.6 | *DLL4* | 3.512 | 1.850144 | Increased |
| ENSG00000168685.14 | *IL7R* | 3.5 | 3.802424 | Increased |
| ENSG00000121858.10 | *TNFSF10* | 3.376 | 3.567363 | Increased |
| ENSG00000163739.4 | *CXCL1* | 3.236 | -0.44897 | Increased |
| ENSG00000137752.22 | *CASP1* | 3.15 | 2.684897 | Increased |
| ENSG00000130775.15 | *THEMIS2* | 3.135 | 3.231943 | Increased |
| ENSG00000185507.19 | *IRF7* | 3.022 | 3.875153 | Increased |
| ENSG00000169429.10 | *CXCL8* | 2.907 | -1.96917 | Increased |
| ENSG00000028277.20 | *POU2F2* | 2.764 | 0.453238 | Increased |
| ENSG00000115604.10 | *IL18R1* | 2.736 | 0.499293 | Increased |
| ENSG00000138378.17 | *STAT4* | 2.695 | 2.145555 | Increased |
| ENSG00000157368.10 | *IL34* | 2.595 | 0.369739 | Increased |
| ENSG00000170298.15 | *LGALS9B* | 2.528 | 2.965212 | Increased |
| ENSG00000136244.11 | *IL6* | 2.503 | -0.23512 | Increased |
| ENSG00000134363.11 | *FST* | 2.481 | 1.895928 | Increased |
| ENSG00000109906.13 | *ZBTB16* | 2.408 | 0.917825 | Increased |
| ENSG00000089041.16 | *P2RX7* | 2.275 | 0.379212 | Increased |
| ENSG00000204632.11 | *HLA-G* | 2.231 | 2.459558 | Increased |
| ENSG00000184371.13 | *CSF1* | 2.22 | 1.125217 | Increased |
| ENSG00000134470.19 | *IL15RA* | 2.14 | 2.119752 | Increased |
| ENSG00000115415.18 | *STAT1* | 2.106 | 2.370024 | Increased |
| ENSG00000115008.5 | *IL1A* | 2.073 | 0.577578 | Increased |
| ENSG00000102524.11 | *TNFSF13B* | 2.069 | 1.73795 | Increased |
| ENSG00000164330.16 | *EBF1* | 2.065 | 2.951442 | Increased |
| ENSG00000123496.7 | *IL13RA2* | 2.024 | 5.897182 | Increased |
| ENSG00000125347.13 | *IRF1* | 2.022 | 1.618451 | Increased |
| ENSG00000134321.11 | *RSAD2* | 10.321 | 9.995906 | Affected |
| ENSG00000169245.5 | *CXCL10* | 10.195 | 8.430441 | Affected |
| ENSG00000107201.9 | *DDX58* | 4.771 | 3.772748 | Affected |
| ENSG00000164342.12 | *TLR3* | 4.216 | 2.7201 | Affected |
| ENSG00000168961.16 | *LGALS9* | 3.756 | 4.338328 | Affected |
| ENSG00000152689.17 | *RASGRP3* | 3.317 | 2.743657 | Affected |
| ENSG00000048052.21 | *HDAC9* | 2.949 | 1.379593 | Affected |
| ENSG00000170581.13 | *STAT2* | 2.748 | 2.681712 | Affected |
| ENSG00000134215.15 | *VAV3* | 2.603 | 2.987545 | Affected |
| ENSG00000169248.12 | *CXCL11* | 4.66 | 4.371531 | Decreased |
| ENSG00000240065.7 | *PSMB9* | 3.567 | 3.780511 | Decreased |
| ENSG00000166923.10 | *GREM1* | 3.214 | -2.05055 | Decreased |
| ENSG00000115602.16 | *IL1RL1* | 2.959 | 0.520912 | Decreased |

### B. Expression of host genes associated with leukocyte proliferation

| ID | Genes in dataset | Expr Log Ratio (UT176 vs UnInf) | Expr Log Ratio (KARP vs UnInf) | Prediction in UT176 (based on measurement direction) |
|---|---|---|---|---|
| ENSG00000171855.6 | *IFNB1* | 8.664 | 8.790328 | Increased |
| ENSG00000164400.5 | *CSF2* | 7.764 | -1.667 | Increased |
| ENSG00000187608.8 | *ISG15* | 5.293 | 5.416531 | Increased |
| ENSG00000108342.12 | *CSF3* | 5.037 | 2.804455 | Increased |
| ENSG00000271503.5 | *CCL5* | 4.801 | 4.102429 | Increased |
| ENSG00000137462.6 | *TLR2* | 4.416 | -0.21786 | Increased |
| ENSG00000172183.14 | *ISG20* | 4.384 | 4.897459 | Increased |
| ENSG00000164342.12 | *TLR3* | 4.216 | 2.7201 | Increased |
| ENSG00000168685.14 | *IL7R* | 3.5 | 3.802424 | Increased |
| ENSG00000172348.14 | *RCAN2* | 2.909 | 1.89052 | Increased |
| ENSG00000100234.11 | *TIMP3* | 2.788 | 1.065608 | Increased |
| ENSG00000028277.20 | *POU2F2* | 2.764 | 0.453238 | Increased |
| ENSG00000138378.17 | *STAT4* | 2.695 | 2.145555 | Increased |
| ENSG00000134215.15 | *VAV3* | 2.603 | 2.987545 | Increased |
| ENSG00000157368.10 | *IL34* | 2.595 | 0.369739 | Increased |
| ENSG00000170091.10 | *NSG2* | 2.557 | 3.988829 | Increased |
| ENSG00000136244.11 | *IL6* | 2.503 | -0.23512 | Increased |
| ENSG00000026950.16 | *BTN3A1* | 2.475 | 2.699843 | Increased |
| ENSG00000170166.5 | *HOXD4* | 2.286 | 2.00549 | Increased |
| ENSG00000089041.16 | *P2RX7* | 2.275 | 0.379212 | Increased |
| ENSG00000184371.13 | *CSF1* | 2.22 | 1.125217 | Increased |
| ENSG00000185950.8 | *IRS2* | 2.142 | -0.59384 | Increased |
| ENSG00000134470.19 | *IL15RA* | 2.14 | 2.119752 | Increased |
| ENSG00000113319.11 | *RASGRF2* | 2.099 | 1.634606 | Increased |
| ENSG00000115008.5 | *IL1A* | 2.073 | 0.577578 | Increased |
| ENSG00000102524.11 | *TNFSF13B* | 2.069 | 1.73795 | Increased |
| ENSG00000114251.13 | *WNT5A* | 4.976 | 3.61382 | Affected |
| ENSG00000162692.10 | *VCAM1* | 4.467 | -0.03961 | Affected |
| ENSG00000168961.16 | *LGALS9* | 3.756 | 4.338328 | Affected |
| ENSG00000152689.17 | *RASGRP3* | 3.317 | 2.743657 | Affected |
| ENSG00000177409.11 | *SAMD9L* | 3.277 | 2.927656 | Affected |
| ENSG00000115602.16 | *IL1RL1* | 2.959 | 0.520912 | Affected |
| ENSG00000023445.13 | *BIRC3* | 2.889 | 1.028695 | Affected |
| ENSG00000163734.4 | *CXCL3* | 2.872 | -0.17656 | Affected |
| ENSG00000152217.16 | *SETBP1* | 2.574 | 1.333599 | Affected |
| ENSG00000140464.19 | *PML* | 2.167 | 1.947424 | Affected |
| ENSG00000169245.5 | *CXCL10* | 10.195 | 8.430441 | Decreased |
| ENSG00000175899.14 | *A2M* | 5.199 | 4.163204 | Decreased |
| ENSG00000131203.12 | *IDO1* | 4.034 | 2.870018 | Decreased |
| ENSG00000057657.14 | *PRDM1* | 3.961 | 1.080978 | Decreased |
| ENSG00000173193.13 | *PARP14* | 3.713 | 3.461054 | Decreased |
| ENSG00000115009.11 | *CCL20* | 3.616 | -1.19333 | Decreased |
| ENSG00000128917.6 | *DLL4* | 3.512 | 1.850144 | Decreased |
| ENSG00000121858.10 | *TNFSF10* | 3.376 | 8.790328 | Decreased |
| ENSG00000169429.10 | *CXCL8* | 2.907 | -1.667 | Decreased |
| ENSG00000118503.14 | *TNFAIP3* | 2.674 | 5.416531 | Decreased |
| ENSG00000170298.15 | *LGALS9B* | 2.528 | 2.804455 | Decreased |
| ENSG00000109906.13 | *ZBTB16* | 2.408 | 4.102429 | Decreased |
| ENSG00000204632.11 | *HLA-G* | 2.231 | -0.21786 | Decreased |
| ENSG00000115415.18 | *STAT1* | 2.106 | 4.897459 | Decreased |

**Supp. Fig. 17. Ot_Karp up-regulates networks associated with (A) organismal growth failure, (B) morbidity and mortality and (C) death.** Connections coloured in red are up-regulated whilst connections in blue are down-regulated.

A. KARP-vs-UnInf UT176-vs-UnInf

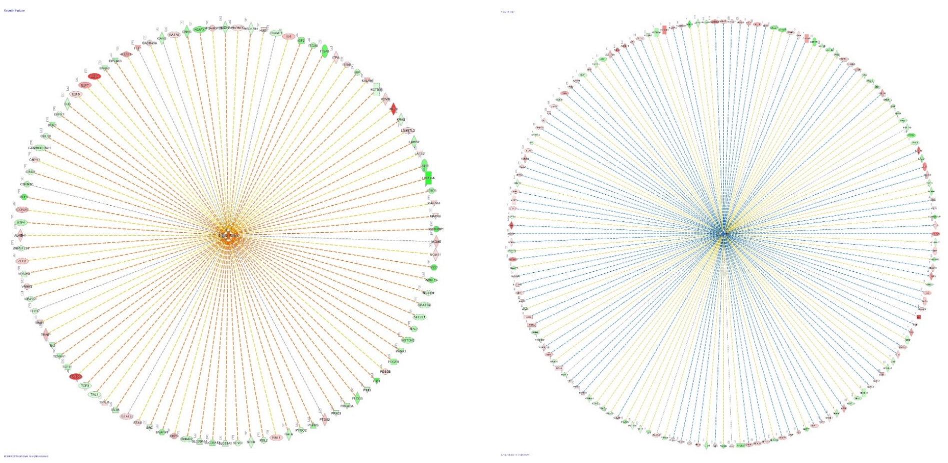

Expression of genes in network associated with Organismal Growth Failure

B. KARP-vs-UnInf UT176-vs-UnInf

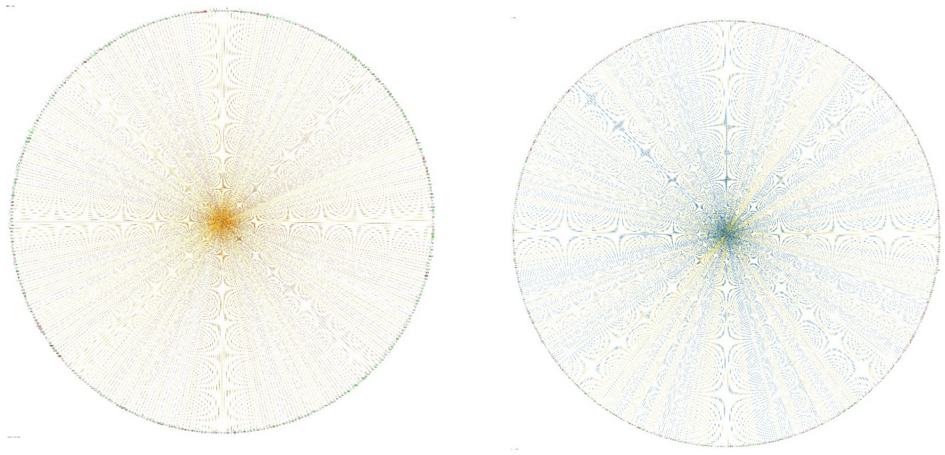

Expression of genes in network associated with Organismal Morbidity and Mortality

C. KARP-vs-UnInf UT176-vs-UnInf

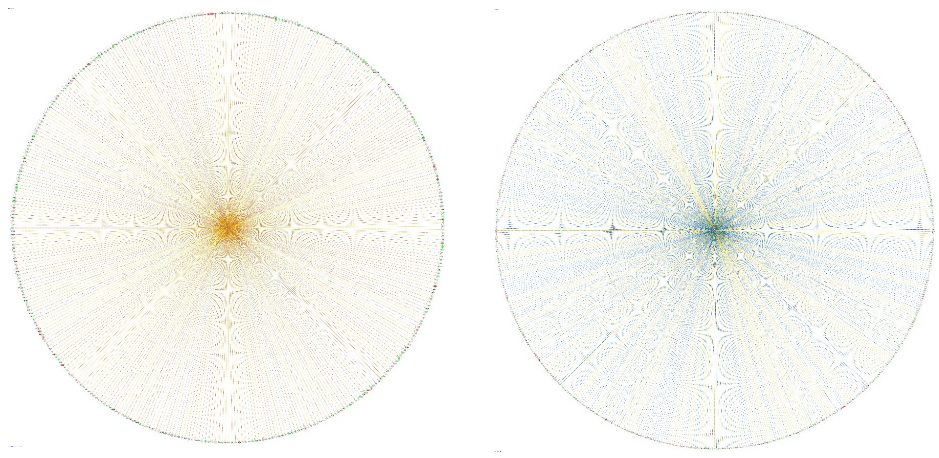

Expression of genes in network associated with Organismal Death

**Supp. Fig. 18. Daily weight measurements of individual mice infected with Ot.**

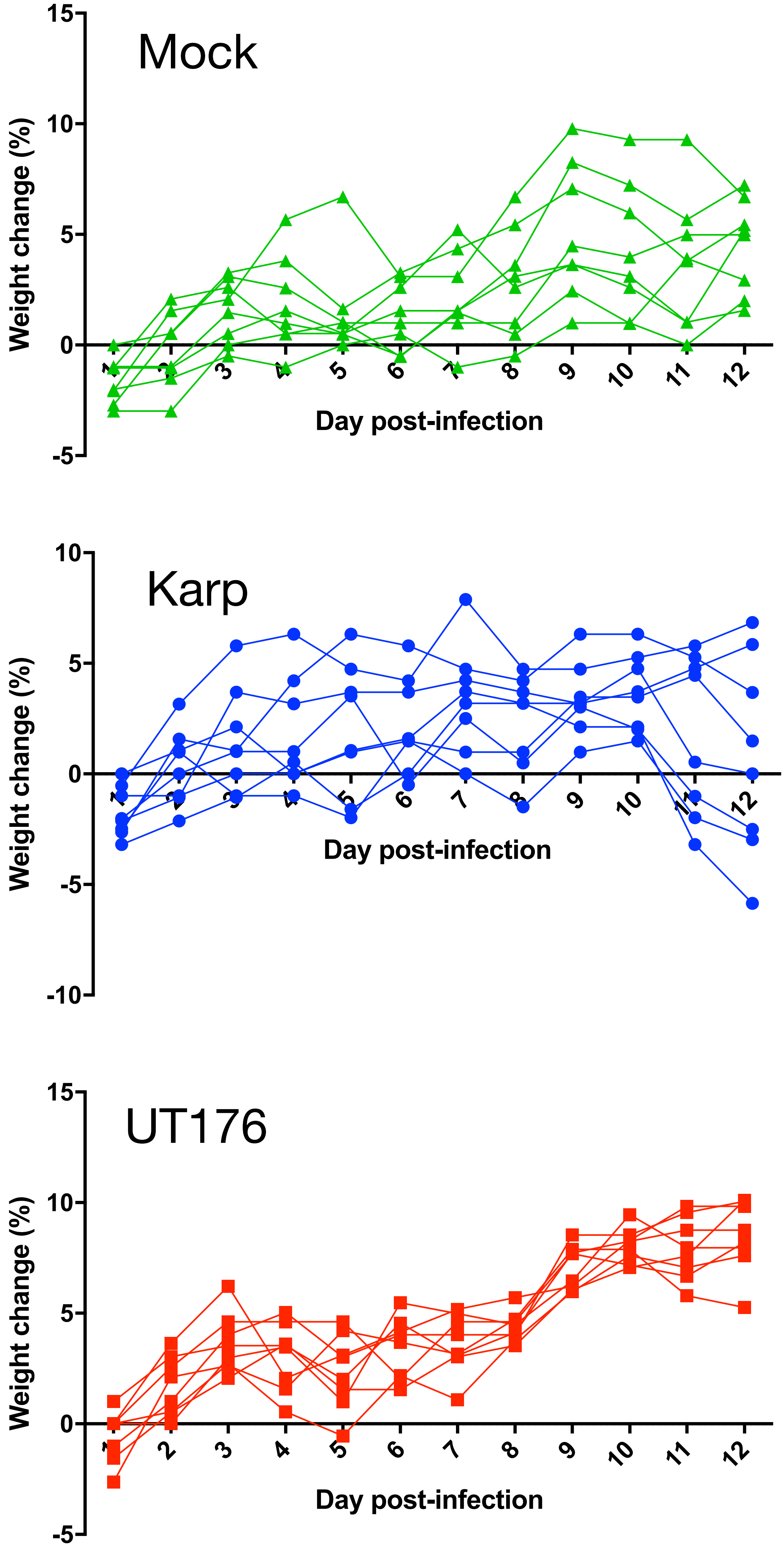

**Supp. Fig. 19. System used for scoring (A) clinical observations and (B) Lesions in H&E stained tissue sections**

A. Mouse clinical observation scoring system

| Parameter | | CODE | SCORE |
|---|---|---|---|
| **Group Appetite** | ate all 6-12 chows | 0 | 0 |
| | ate 1-5 chows | 1 | 1 |
| | ate 0 to 1/2 chow | 2 | 2 |
| **Activity** | move around | 0 | 0 |
| | locomotion after slight stimulation | A1 | 1 |
| | move slowly after moderate stimulation | A2 | 2 |
| | unable to move | A3 | 3 |
| **Hair coat** | well-groomed hair coat | 0 | 0 |
| | rough hair coat | R1 | 1 |
| | ungroomed, very rough hair coat, and dirty | R2 | 2 |
| **Total score** | 7 | | |

B.

Lesion scoring system

0 = Normal tissue
1 = Minimal lesion severity and extent
2 = Mild
3 = Moderate
4 = Marked
5 = Severe

**Supp. Fig. 20. Histopathological analysis of Karp- and UT176-infected mouse tissues.** A-D. Karp-infected mouse lungs show increased cellular infiltration, compared with UT176 infected counterpart. E-I. Karp-infected mouse liver shows increased necrosis and inflammation, compared with UT176 infected counterpart. Arrows in E&I: boundaries of necrotic (nec) zone; Arrows in F&J: bigger and multiple inflammatory foci with cellular aggregation; Arrows in H: smaller and fewer inflammatory foci with cellular aggregation; Arrows in K: perivascular inflammatory foci. Figs I-L are magnified section of E-H, respectively.

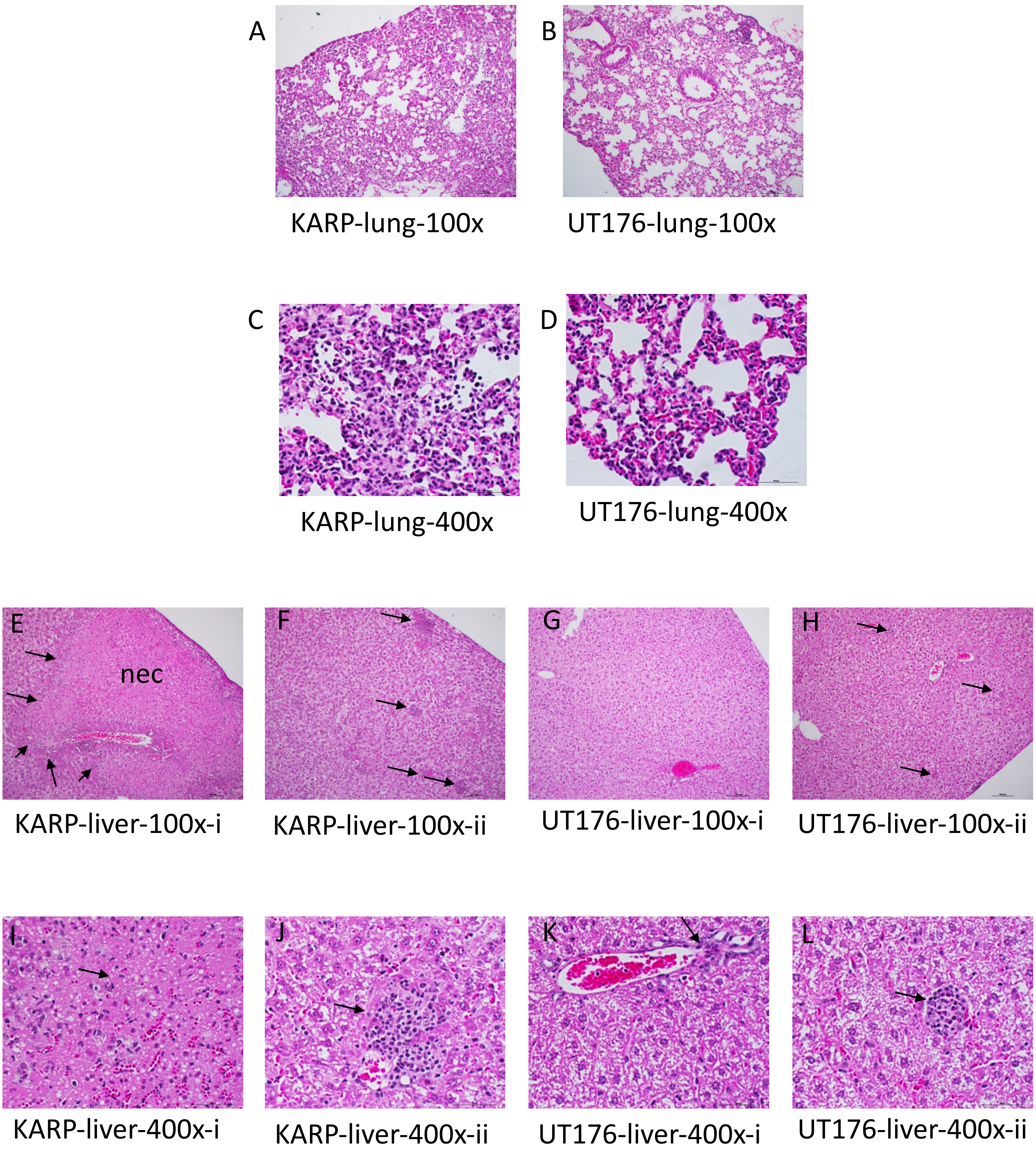
