## Supplementary text for "Dual RNA-seq provides insight into the biology of the neglected intracellular human pathogen *Orientia tsutsugamushi*"

Bozena Mika-Gospodorz[#a], Suparat Giengkam[#b], Alexander J. Westermann[a,c], Jantana Wongsantichon[b], Willow Kion-Crosby[d], Suthida Chuenklin[b], Loo Chien Wang[e], Piyanate Sunyakumthorn[f], Radoslaw M. Sobota[e], Selvakumar Subbian[h], Jörg Vogel[a,c], Lars Barquist[#*a,g] and Jeanne Salje[#*b,h,i]

Running title: RNA sequencing of *Orientia tsutsugamushi*

Helmholtz Institute for RNA-based Infection Research (HIRI), Helmholtz Centre for Infection Research (HZI), Würzburg, Germany[a]

Mahidol-Oxford Tropical Medicine Research Unit, Faculty of Tropical Medicine, Mahidol University, Bangkok, Thailand[b];

Institute for Molecular Infection Biology (IMIB), University of Würzburg, Würzburg, Germany[c]

Rutgers, the State Univeristy of New Jersey, New Jersey, USA[d]

Functional Proteomics Laboratory, Institute of Molecular and Cell Biology, Agency for

Science, Technology and Research (A*STAR), Singapore[e].

Armed Forces Research Institute of Medical Sciences, Bangkok, Thailand[f];

Faculty of Medicine, University of Würzburg, Würzburg, Germany[g];

Public Health Research Institute, Rutgers University, New Jersey, USA[h];

Centre for Tropical Medicine and Global Health, Nuffield Department of Medicine, University of Oxford, Oxford, United Kingdom[i];

SingMass - National Mass Spectrometry Laboratory, Institute of Molecular and Cell Biology, Agency for Science, Technology and Research (A*STAR), Singapore[j]

[#] these authors contributed equally

**Codon bias in Ot**

A comparison of our RNAseq and proteomics datasets revealed a further 218 core genes detected by RNAseq but not proteomics. Some of this discrepancy is likely due to differences in sample preparation and sensitivities between the two methods, loss of secreted proteins, and differences in protein stability. However, some of this difference might also be explained by differences in codon usage resulting in differences translation efficiency. To address this question, we analyzed global codon usage in Ot and compared it with two other alpha-proteobacterial species, *Caulobacter crescentus* and *Rickettsia typhi* (Fig. 4A), and the gamma-proteobacterium *E. coli*. Genome-wide biases in Ot and *R. typhi* were very similar, but they differed from those in *C. crescentus* and *E. coli,* supporting previous analyses suggesting that codon bias in obligate intracellular bacteria is not strong, which may reflect slow growth rates not limited by the speed of translation[1–3]. We observed no clear correlation between codon usage and the presence or absence of a cognate tRNA in Ot. Ot encodes 34 out of 61 possible amino-acid coding tRNAs, leaving 27 potential codons without a cognate tRNA molecule. Of the 20 amino acids, 7 have a strong bias against the cognate Ot tRNA, 10 have biases for both cognate and non-cognate Ot tRNAs and only 3 have biases for the cognate Ot tRNA (Supp. Fig. 8-9), unlike codon biases in other organisms, which correlate with tRNA copies[4–7]. There may be mechanisms by which bias towards non-cognate tRNA is preferential, or perhaps fast translation rates are not strongly selected for in Ot.

To determine whether Ot uses codon bias to regulate gene expression we compared codon usage between different groups of Ot genes (Supp. Fig. 8B, C). When comparing core and non-core gene groups, seventeen out of twenty amino acids showed differences in codon usage (Supp. Fig. 8-9), likely reflecting the horizontal acquisition of RAGE genes and possible differences in their subsequent optimization of translation.

We compared codon usage between core genes that were either detected or not detected in our proteomics dataset and found nine amino acids that were differentially encoded between these groups (Supp. Fig. 8B, C). The differences were statistically significant but smaller than those seen in other organisms, such as *E. coli*[8,9]. We compared these codon usage biases to those of a group of seven genes whose expression was high based on relative peptide levels in the proteomics dataset (RpoB1, RpoB2, GroEL, DnaK, GdhA, HtpG, TSA56) and observed the same trend as seen between those not detected and detected in the proteomics dataset. Thus, these differences likely reflect a true relationship between codon usage and translation, although the contribution of this bias to protein expression remains unknown.
